# Developmental heterogeneity of microglia and brain myeloid cells revealed by deep single-cell RNA sequencing

**DOI:** 10.1101/406363

**Authors:** Qingyun Li, Zuolin Cheng, Lu Zhou, Spyros Darmanis, Norma Neff, Jennifer Okamoto, Gunsagar Gulati, Mariko L. Bennett, Lu O. Sun, Laura E. Clarke, Julia Marschallinger, Guoqiang Yu, Stephen R. Quake, Tony Wyss-Coray, Ben A. Barres

## Abstract

Microglia are increasingly recognized for their major contributions during brain development and neurodegenerative disease. It is currently unknown if these functions are carried out by subsets of microglia during different stages of development and adulthood or within specific brain regions. Here, we performed deep single-cell RNA sequencing (scRNA-seq) of microglia and related myeloid cells sorted from various regions of embryonic, postnatal, and adult mouse brains. We found that the majority of adult microglia with homeostatic signatures are remarkably similar in transcriptomes, regardless of brain region. By contrast, postnatal microglia represent a more heterogeneous population. We discovered that postnatal white matter-associated microglia (WAM) are strikingly different from microglia in other regions and express genes enriched in degenerative disease-associated microglia. These postnatal WAM have distinct amoeboid morphology, are metabolically active, and phagocytose newly formed oligodendrocytes. This scRNA-seq atlas will be a valuable resource for dissecting innate immune functions in health and disease.

**Highlights:**

- Myeloid scRNA-seq atlas across brain regions and developmental stages
- Limited transcriptomic heterogeneity of homeostatic microglia in the adult brain
- Phase-specific gene sets of proliferating microglia along cell cycle pseudotime
- Phagocytic postnatal white matter-associated microglia sharing DAM gene signatures

## Introduction

Microglia are brain parenchymal macrophages that are implicated in numerous neurological diseases, such as Alzheimer’s disease, amyotrophic lateral sclerosis, stroke, and brain tumors (Colonna and Butovsky, 2017; Prinz et al., 2011). In addition to their classical immune surveillance and scavenging functions, microglia have recently been found to actively participate in neural development by modulating neurogenesis and pruning synapses (Cunningham et al., 2013; Li and Barres, 2017; Paolicelli et al., 2011; Schafer et al., 2012; Ueno et al., 2013). Despite the importance of these multitasking cells, little is known about their molecular heterogeneity under physiological conditions and especially during development when they perform many critical non-immune functions. In addition, due to their transcriptomic resemblance to other myeloid cells which may infiltrate the brain parenchyma in disease (Goldmann et al., 2016; Prinz et al., 2017), a systematic comparison between microglia and these related immune cells remains an imperative task. Therefore, identifying functional subsets of microglia in the context of other myeloid cells is an essential step towards a better understanding of brain development as well as guiding therapeutic interventions for disease and injury.

Microglia and most other tissue macrophages are long-lived, self-renewing cells that are generated by waves of erythro-myeloid progenitors in the yolk sac, independently of bone marrow-derived cells (Gomez Perdiguero et al., 2015; Hoeffel et al., 2015; Li and Barres, 2017). In mice, microglia migrate to the brain around embryonic day 9.5 (E9.5) and blood-brain barrier closure around E13.5 has been proposed as a mechanism to confine microglia inside the parenchyma (Ginhoux et al., 2010). Consistent with this convoluted developmental route, bulk RNA-sequencing (RNA-seq) data demonstrated a roughly step-wise differentiation program for microglia (Matcovitch-Natan et al., 2016). However, the reliance on general surface markers in these studies could overlook microglial heterogeneity, particularly potential transient populations during development, thereby underestimating developmental complexity of microglia. Furthermore, although mature microglia in different brain regions were shown to have uneven distribution with distinct morphologies (Lawson et al., 1990), which seem to correlate with region-specific expression profiles (Ayata et al., 2018; De Biase et al., 2017; Grabert et al., 2016), it remains unclear whether there are molecularly defined subtypes of microglia in the adult brain and, if so, how they are distributed across brain regions.

We took an unbiased approach to investigate the heterogeneity of microglia along with other brain myeloid cells by performing deep single cell RNA-seq (scRNA-seq) on sorted cells across mouse brain regions and developmental stages. scRNA-seq has been proved as a powerful tool for dissecting cellular diversity from complex organs with minimal prior knowledge (Papalexi and Satija, 2018). We utilized the Smart-seq2 approach on sorted cells, due to its superior sensitivity and remarkable accuracy (Svensson et al., 2017; Ziegenhain et al., 2017). In total, we sequenced 1922 cells to over 1 million raw reads per cell. Clustering analysis of this complex dataset identified 15 distinct cell populations, and differential gene expression analysis of these clusters further generated lists of marker genes for each population. Two microglia clusters expressed signature genes for dividing cells, which we used to reconstruct cell cycle phases and produced phase-specific gene sets for microglia. We also found that postnatal and adult choroid plexus macrophages were separated into distinct clusters, suggesting a developmental phenotypic switch for these particular brain resident macrophages. We created a searchable web interface for this dataset as an integrated component of our widely used brain RNA-seq website: www.brainrnaseq.org.

Surprisingly, we found little population-wise heterogeneity among adult homeostatic microglia at the whole transcriptomic level. Furthermore, scRNA-seq and bulk RNA-seq consistently demonstrated that the typical homeostatic microglia, representing the vast majority of adult microglia in different brain regions, were remarkably similar in global gene expression despite tissue origins. By contrast, we observed much higher heterogeneity in early postnatal microglia. Particularly, we identified a population of developing white matter-associated microglia (WAM), that shared transcriptional signatures with degenerative disease-associated microglia (DAM) (Keren-Shaul et al., 2017; Krasemann et al., 2017). We characterized specific molecular markers for postnatal WAM, and we further showed that they were amoeboid, metabolically active and highly phagocytic. Postnatal WAM transiently appeared in developing corpus callosum and cerebellar white matter around the first postnatal week, when they mainly engulfed newly formed oligodendrocytes. Interestingly, unlike DAM (Keren-Shaul et al., 2017; Krasemann et al., 2017), appearance of postnatal WAM did not depend on a TREM2-APOE axis, suggesting that different signals may trigger the emergence of these two microglial populations.

We believe that our myeloid cell atlas with unprecedented resolution in gene expression across developmental stages and anatomical domains will serve as a valuable tool for dissecting innate immune cell contributions to normal development and brain disease.

## Results

### Clustering of brain myeloid cells across developmental stages by deep scRNA-seq

To understand the spatiotemporal heterogeneity of microglia and other brain myeloid cells in an unbiased manner, we used a semi-automated Smart-seq2 platform to perform single-cell RNA sequencing (scRNA-seq) on index sorted innate immune cells from the mouse brain (Figure 1A). We sequenced a total of 1922 cells to a depth of at least 1 million raw reads per cell, from embryonic day 14.5 (E14.5), postnatal day 7 (P7) and adult (P60) brain tissues. We reasoned that deeper sequencing would allow us to detect more genes compared with other scRNA-seq technologies, which could reveal even subtle differences among possible microglial subpopulations. We also analyzed non-microglia myeloid cells, due to their resemblance to parenchymal microglia and possible links to degenerative diseases and injury. To uncover microglial regional heterogeneity, we isolated myeloid cells (for P7 and P60 stages) from six brain regions: cortex, cerebellum, hippocampus, striatum, olfactory bulb, and choroid plexus. We used a recently developed protocol for cell extraction to minimize microglial activation (Bennett et al., 2016). For the E14.5 stage, we gated on c-Kit-CD45^+^; for the P7 and P60 stages, we gated on CD45^low^CD11b^+^ versus CD45^hi^CD11b^+^ for microglia and myeloid cells, respectively (Figures 1B and S1A). Fluorescence intensities from other relevant markers, such as F4/80 (macrophage marker), CD41 (hematopoietic progenitor marker) and TMEM119 (homeostatic microglia marker), were recorded as part of the metadata in this multidimensional dataset (Figures 1A and S1A).

**Figure 1.**
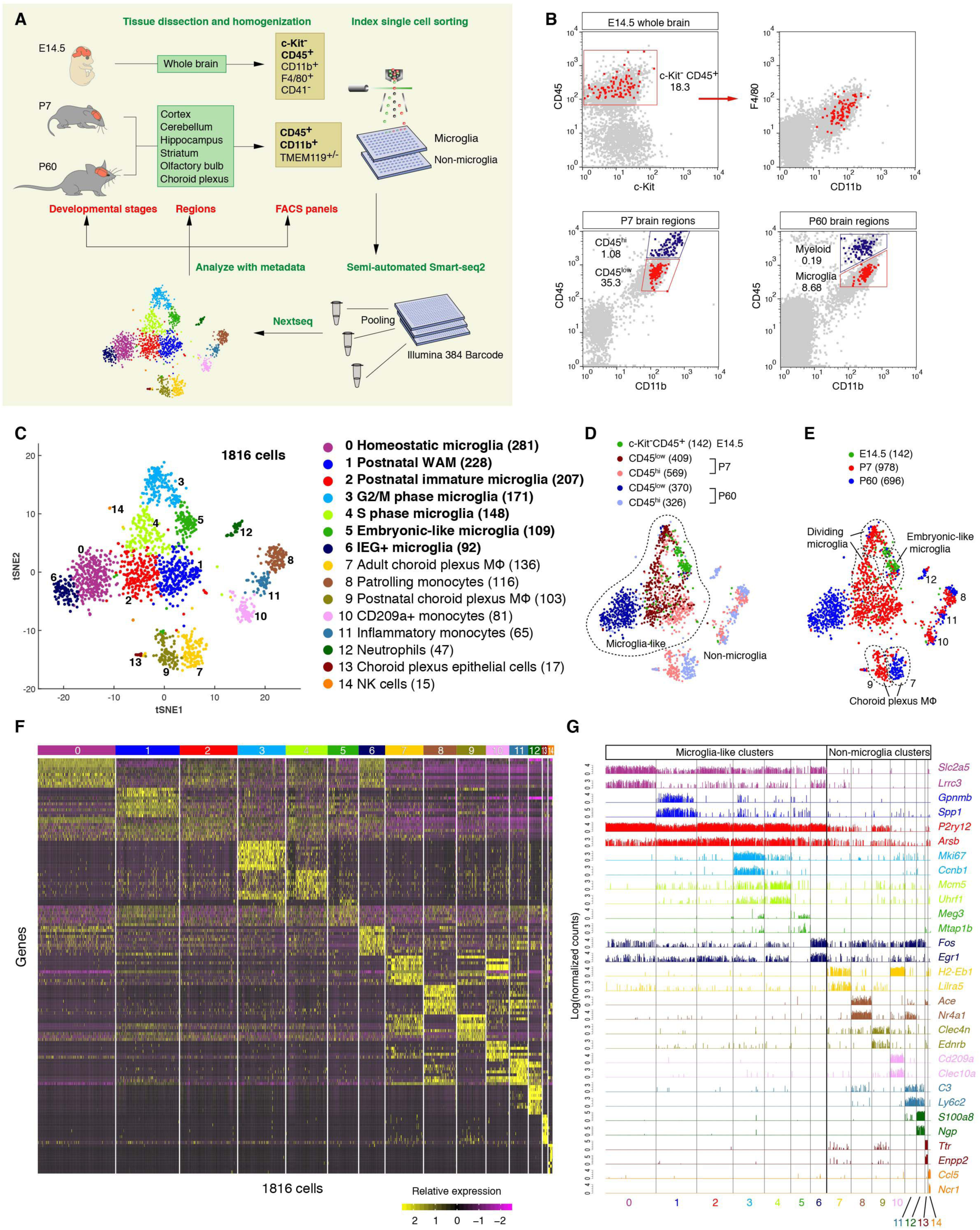
Clustering of brain myeloid cells across developmental stages by deep scRNA-seq. (A) Schematic graph showing the experimental design for isolating microglia and other brain myeloid cells from different brain regions and across developmental stages. Whole brains were used for E14.5 samples, and c-Kit-CD45^+^ (bold) myeloid cells were gated for single-cell index sorting. These cells were also CD11b^+^F4/80^+^CD41-, which were recorded as metadata. For P7 and P60 samples, tissues from named six brain regions were dissected, and cells were gated on CD45^+^CD11b^+^ (bold) for microglia (CD45^low^, also TMEM119^+^), or other myeloid cells (CD45^hi^, also TMEM119-). Libraries were made by a semi-automated Smart-seq2 protocol, and every 320-380 samples (each with a unique barcode) were pooled for Illumina sequencing. scRNA-seq data were analyzed together with all metadata. (B) Representative FACS plots showing the cells sequenced (colored in red or blue) in our study. Microglia and myeloid cells were sorted separately. (C) tSNE plot showing 15 clusters from all 1816 cells (passed QC) identified by the clustering algorithm. Cell population annotations are labeled on the right with all microglia-like clusters in bold. The number of cells in each cluster is given in parentheses. (D) Overlaying FACS gating information onto the same tSNE plot in (C) highlighting distinctions between microglia-like and non-microglia clusters. The vast majority of P60 microglia are CD45^low^, while P7 microglia are comprised of both CD45^low^ and CD45^hi^ cells. Almost all cells from non-microglia clusters are CD45^hi^. (E) Overlaying developmental stage information onto the same tSNE plot in (C) showing the shift of microglia-like clusters from embryonic-like cells (green) to postnatal (red) and then to adult microglia (blue). Both P7 (red) and P60 (blue) cells are found in each of monocyte (cluster 8, 10, 11), or neutrophil (cluster 12) clusters, whereas choroid plexus macrophages are separated into two clusters (cluster 7 and 9) largely according to their developmental stages. Two dividing microglia clusters are also circled, which include mainly P7 and E14.5 cells with some P60 cells (blue). (F) Heatmap showing the top 20 markers (or all markers if less than 20) for each of the 15 clusters which are distinguished by unique sets of genes. (G) Bar plots showing gene expression levels of two representative markers for each cluster across all 1816 cells. See also Figure S1, Table S1.

Using stringent criteria, 1816 cells (94.5% of 1922) passed quality control for downstream analysis, including 142 E14.5 cells, 978 P7 cells, and 696 P60 cells from various brain regions (Figures S1B-S1D; see the Methods). To assess the sensitivity and accuracy of the data, we analyzed the sequencing results for External RNA Controls Consortium (ERCC) Spike-In RNA, and found that our Smart-seq2 platform had nearly single molecule detection limit and high precision (R=0.98) (Figures S1E and S1F; see the Methods). When comparing our sequencing data for adult microglia with two other published scRNA-seq datasets (Keren-Shaul et al., 2017; Matcovitch-Natan et al., 2016), our study detected about 3 times as many genes per cell, and produced higher detection rates for over 80% of genes (Figures S1G and S1H). In summary, we generated a high-quality multidimensional scRNA-seq dataset for region-specific microglia and brain myeloid cells across developmental stages, and we incorporated these data into our widely used brain RNA-seq website (www.brainrnaseq.org).

To study the transcriptomic heterogeneity of brain myeloid cells, we carried out unsupervised clustering analysis for all 1816 cells, which gave rise to 15 distinct clusters (Figure 1C). Through differential gene expression analysis, we discovered a list of enriched genes for each cluster as potential markers (Table S1). Expression of these genes uniquely or in combinations represented individual cluster identities (Figures 1F and 1G). Based on these gene lists and other metadata, such as the developmental stage and FACS gating information for the cells from each cluster (Figures 1D and 1E), we annotated the 15 clusters into distinct cell types or states. This included 7 microglia clusters, 2 choroid plexus macrophage clusters, 3 monocyte clusters and 3 other immune or epithelial cell clusters (Figure 1C). Of note, some clusters had less than 20 cells (e.g. cluster 14, Natural killer cells), but were robustly identified as distinct and biologically meaningful clusters, suggesting the reliability of the current technology even for small numbers of cells.

Interestingly, we observed that different clusters of microglia, choroid plexus cells or monocytes tended to be most closely juxtaposed on the tSNE plot, implicating the overall similarities between clusters of each cell type (Figure 1C). For example, neighboring clusters 0-6 were deemed “microglia-like” cells (Figures 1C and 1D). They all expressed microglial signature genes such as *P2ry12* and *Slc2a5* (Figure 1G) (Bennett et al., 2016; Zhang et al., 2014), and these cells were largely isolated from the CD45^low^ gate, whereas cells from other “non-microglia” clusters were almost exclusively from the CD45^hi^ gate (Figure 1D). We noticed the presence of many CD45^hi^ cells from the P7 stage in “microglia-like” clusters, suggesting the heterogeneity of CD45 immunophenotypes in early postnatal microglia. Furthermore, according to the developmental stages, these 7 microglia clusters were segregated into unidirectionally shifted domains on the tSNE plot, consistent with progressive changes in gene expression during microglial development (Figure 1E) (Matcovitch-Natan et al., 2016).

### Postnatal transcriptional alteration of choroid plexus macrophages

We first focused on non-microglia clusters. Consistent with the bone marrow origin of circulating myeloid cells, we observed that monocyte and neutrophil clusters comprised intermingled P7 and P60 cells within each cluster (Figures 1C and 1E). On the other hand, choroid plexus macrophages (CP MΦ) as resident macrophages at central nervous system (CNS) interfaces, have yolk-sac and bone marrow dual origins (Goldmann et al., 2016). Remarkably, CP MΦ were separated into two distinct clusters (cluster 7 and cluster 9) largely confined to their developmental stages (Figure 1E). Only in the adult CP MΦ cluster (cluster 7) did we detect high levels of MHC-II gene expression (e.g. *H2-Ab1, Cd74, H2-Aa, H2-Eb1*) (Figure 2A). Along with these genes involved in antigen presentation, we also found elevated expression of other immunoregulatory genes, such as *Fkbp5, Tsc22d3, Axl* and *Lilra5* in cluster 7. By contrast, postnatal CP MΦ (cluster 9) expressed many genes that function in endocytosis and intracellular trafficking of nutrients or signaling molecules (e.g. *Snx6, Snx2, Dab2, Cd36, Ap1b1, Mrc1*) (Figure 2A). We identified *H2-Eb1* and *Lilra5* as molecular markers for adult CP MΦ and *Clec4n* for their postnatal counterparts, and validated this selective gene expression by RNA *in situ* and immunohistochemistry (Figures 2B-2D). Our lab has previously demonstrated that microglia turn on homeostatic genes during postnatal development (Bennett et al., 2016). Here we show that CP MΦ are a second example of brain resident macrophages that undergo a postnatal transcriptional alteration.

**Figure 2.**
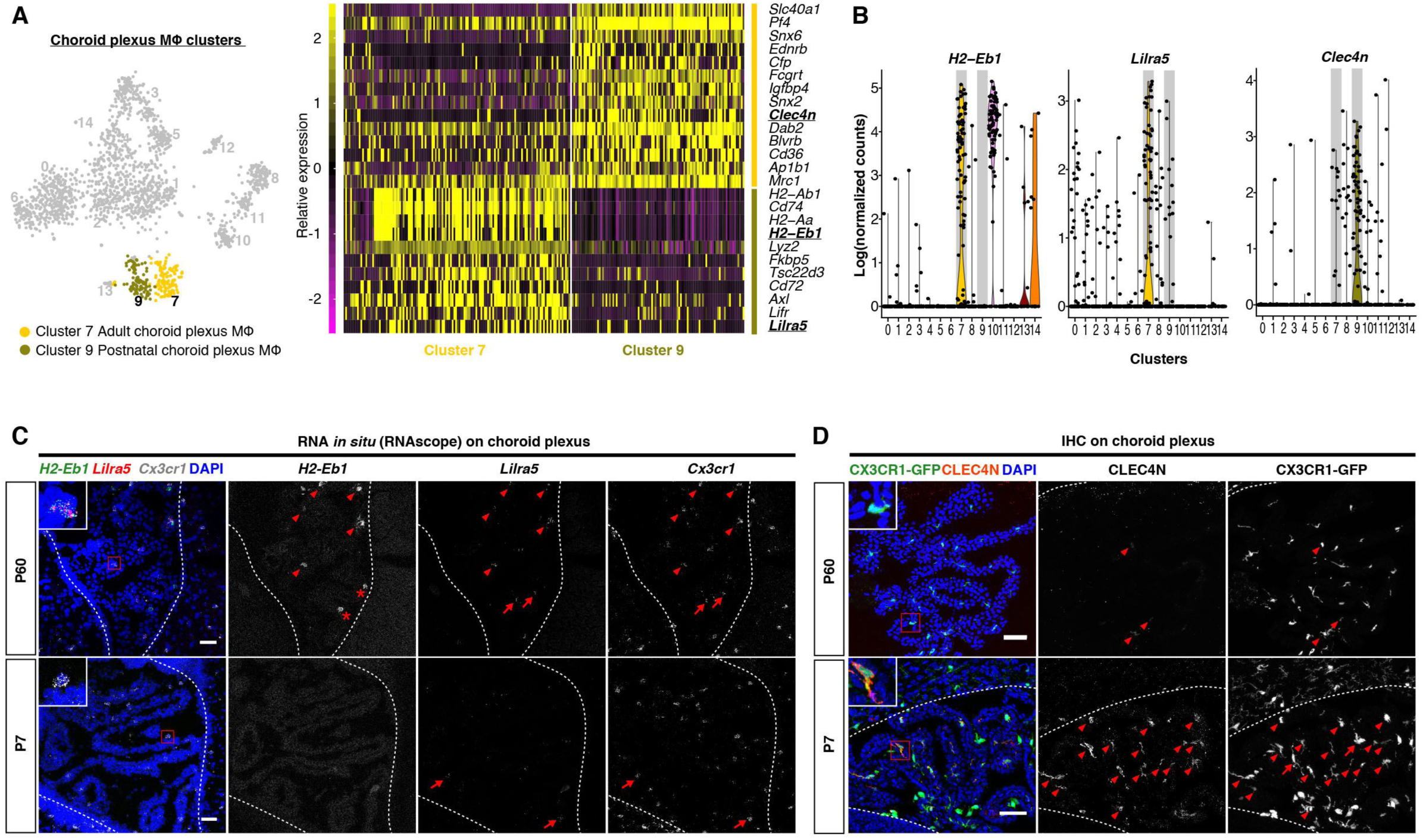
Postnatal transcriptional alteration of choroid plexus macrophages (CP MΦ). (A) tSNE plot on the left highlighting two stage-specific CP MΦ clusters. Heatmap on the right showing differentially expressed genes between two CP MΦ clusters. Underlined genes (bold) were validated by histology in (C) and (D). (B) Violin plots showing expression levels of three marker genes for CP MΦ across 15 clusters identified in Figure 1C. The relevant cluster 7 and 9 are shaded in gray. *H2-Eb1* and *Lilra5* are highly expressed by adult CP MΦ, whereas *Clec4n* is highly enriched in postnatal CP MΦ. (C) Validation of abundant *H2-Eb1* and *Lilra5* expression in P60 CP MΦ but not in P7 cells by RNA *in situ*. Arrow heads point to *H2-Eb1*/*Lilra5*/*Cx3cr1* triple positive CP MΦ in P60 tissue. *H2-Eb1*-*Cx3cr1*^+^*Lilra5*^+^ (arrows) cells were occasionally seen in both stages. *H2-Eb1*^+^*Lilra5*-*Cx3cr1*-non-myeloid cells are labeled with asterisks. (D) Validation of abundant CLEC4N expression in P7 CP MΦ but not in P60 cells by immunostaining. Arrow heads point to CLEC4N/CX3CR1-GFP double positive cells, which are more prominent in P7. CLEC4N-CX3CR1-GFP^+^ (arrow) cells are also present in P7. Scale bars in (C) and (D) are 50um.

### Limited transcriptomic heterogeneity of adult homeostatic microglia

Within the 7 microglia clusters, adult microglia were almost exclusively present in cluster 0 and cluster 6 (Figures 1C-1E). Both clusters expressed similar levels of homeostatic microglial signature genes (Figure S2A), with the exception of slightly reduced *P2ry12* expression in cluster 6 (Figure 3A). Compared with cluster 0, only 27 total genes (including *P2ry12*) were differentially expressed in cluster 6, out of which half were immediate early genes (IEG) (Table S2; Figure 3A). Therefore, we annotated cluster 6 as “IEG+” microglia. Since these immediate early genes can be rapidly induced in response to environmental stimuli, and the microglia extraction procedure is likely to trigger such an artificial response, we examined the expression of the two most up-regulated IEG genes, *Fos* and *Egr1*, on tissue sections. Indeed, adult microglia stained negative for *Fos*/*Egr1* transcripts or proteins, whereas their expression was readily detected in the surrounding non-microglial cells (Figures 3B, 3C, S2B and S2C). This result is consistent with recent studies showing that immediate early gene expression by microglia was only found in sorted cells but not with the RiboTag technology (Ayata et al., 2018; Haimon et al., 2018). Taken together, if we consider the contribution of IEGs to the clustering result as an experimental artifact, the vast majority of adult microglia from different brain regions appear to show the classical homeostatic phenotype with remarkable transcriptomic similarity.

**Figure 3.**
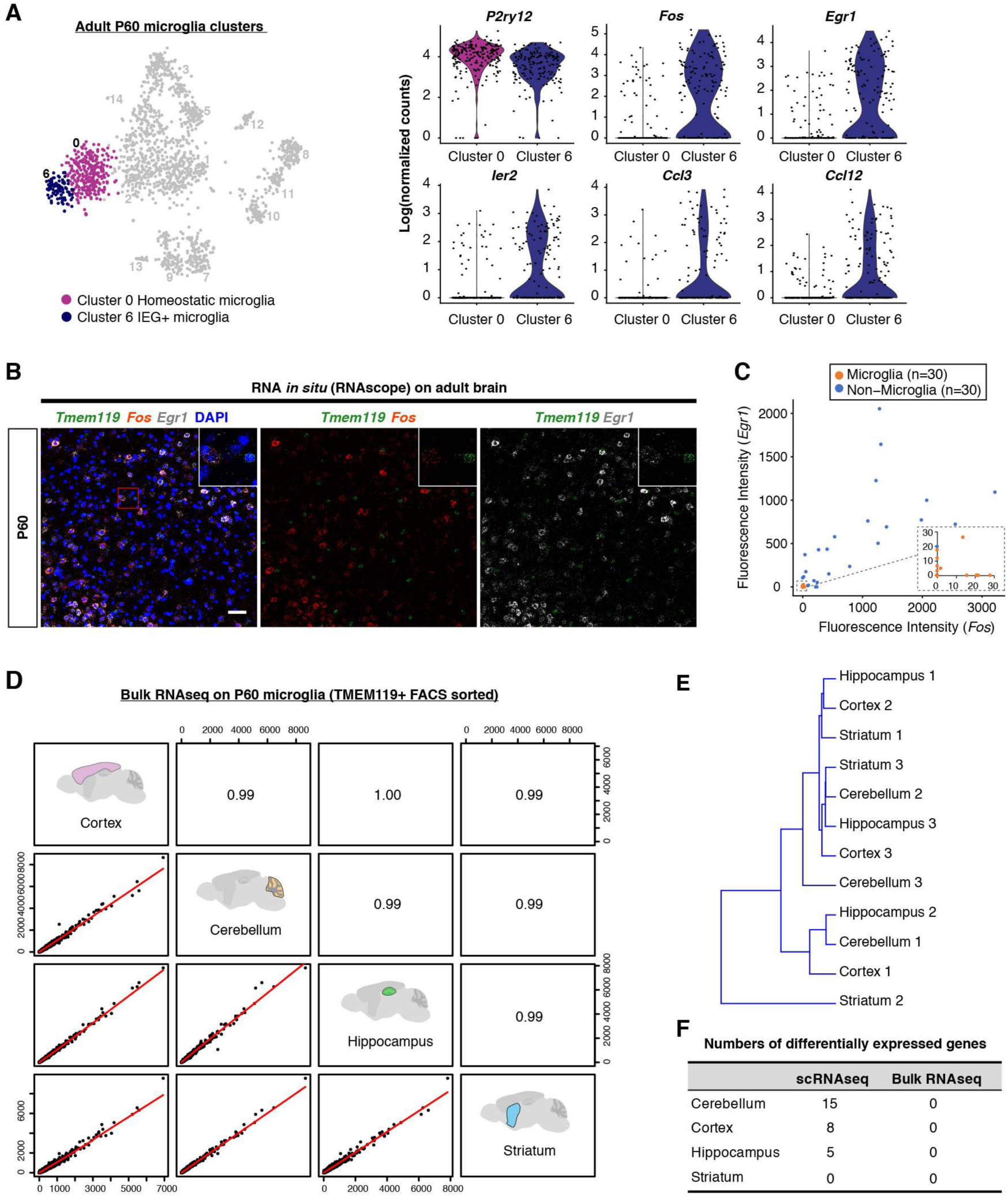
Limited transcriptomic heterogeneity of adult homeostatic microglia across brain regions. (A) tSNE plot on the left highlighting two P60 microglia clusters. Violin plots on the right showing examples of differentially expressed genes between two P60 microglia clusters. (B) RNA *in situ* on fresh frozen sections of adult brains showing negative expression of *Fos* and *Egr1* in microglia (*Tmem119*^+^), and positive signals from surrounding non-microglia cells. (C) Quantification of fluorescence signals in (B). n=30 cells each, for microglia and non-microglia cells. (D) Pearson correlation between FACS sorted bulk RNA-seq samples from different brain regions showing resemblance of region-specific homeostatic (TMEM119+) microglia at the transcriptomic level in the adult stage. (E) Dendrogram showing hierarchical clustering of bulk RNA-seq samples. Replicates from each brain region do not cluster according to tissue origins. (F) Numbers of differentially expressed genes between homeostatic microglia from different regions by scRNA-seq and bulk RNA-seq. Scale bar in (B) is 50um. See also Figure S2, Table S2, Table S3.

Since we collected cells from different areas of the brain, these analyses imply that adult homeostatic microglia have a similar expression profile despite varying regions-of-origin. Indeed, we found that less than 20 genes were differentially expressed (FDR < 0.05) in typical adult microglia (cluster 0) between regions (Figure 3F; Table S3). To rule out a sensitivity issue of scRNA-seq, we performed region-specific bulk RNA-seq of highly purified homeostatic microglia sorted with TMEM119 antibodies, which label over 90% of microglia in the adult brain (Bennett et al., 2016). Consistent with our scRNA-seq data, samples from cortex, cerebellum, hippocampus and striatum were highly correlated (R>0.99), and individual samples did not cluster according to tissue origins, suggesting remarkable similarities between homeostatic microglia from different brain regions (Figures 3D and 3E). Moreover, we could not detect any differentially expressed genes (FDR < 0.05) between regions from the bulk samples (Figure 3F; Table S3). Conversely, we found that certain genes previously attributed to microglial regional heterogeneity were largely expressed by different populations of non-microglial cells from our scRNA-seq data (Grabert et al., 2016) (Figure S2D). These data suggest that classical adult microglia with homeostatic signatures (e.g. *Tmem119* and *P2ry12*), as the most dominant microglial population in the healthy brain, have only limited, if any, transcriptomic heterogeneity across brain regions.

### Phase-specific gene expression of dividing microglia along cell cycle pseudotime

We next characterized postnatal microglia, which were distributed among clusters 0-5 (two P7 cells in the IEG+ cluster 6 were excluded from the analysis) (Figure 4A, upper left). These included two clusters corresponding to dividing cells in two cell cycle phases (cluster 3 and 4), one embryonic-like cluster (cluster 5), two postnatal cell-specific clusters (cluster 1 and 2), and a few cells in the adult homeostatic cluster (cluster 0) bordering cluster 2.

**Figure 4.**
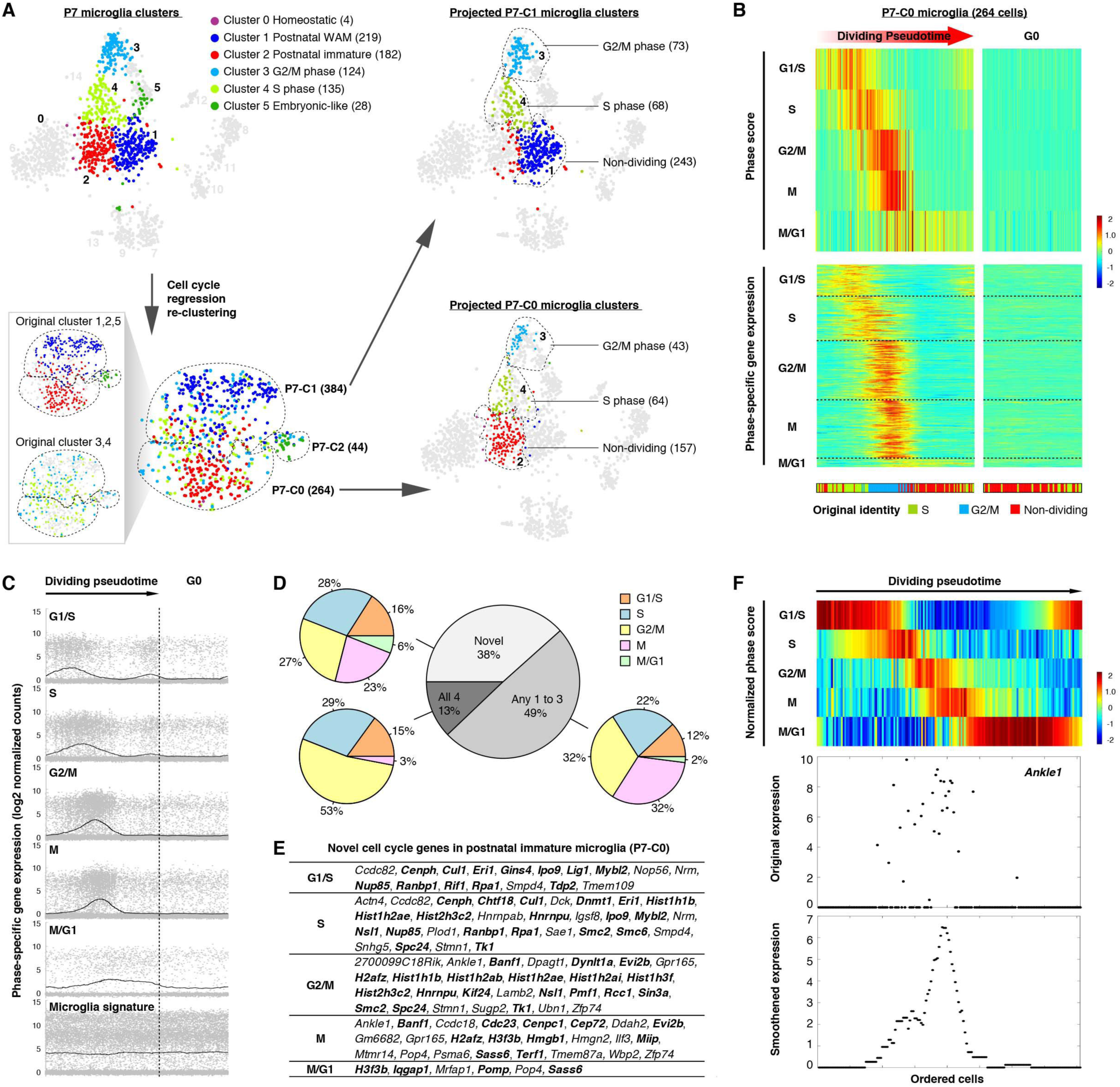
Phase-specific gene expression of dividing microglia (P7-C0 cluster) along cell cycle pseudotime. (A) tSNE plots demonstrating cell cycle regression and re-clustering of P7 microglia. tSNE plot on the upper left highlighted P7 microglia and their cluster identities as determined in Figure 1C. tSNE plot on the lower left shows the result after cell cycle regression and re-clustering. Two tSNE plots in the box highlight the original cluster identities in order to demonstrate consistency of two analyses (before and after cell cycle regression) and how the original dividing clusters (cluster 3 and 4) are distributed into P7-C0, P7-C1 and P7-C2 following cell cycle regression. tSNE plots on the upper right and lower right highlight P7-C1 and P7-C0 microglia in the original plot, respectively, based on the re-clustering analysis. Cells in all tSNE plots are color coded exactly the same way as in Figure 1C. Numbers of cells in each cluster are given in parentheses. (B) – (F) are detailed analysis for the 264 P7-C0 microglia identified in (A). Heatmap (upper panel) showing pseudotime ordering of P7-C0 microglia based on raw phase scores. Each column is a cell, and each row denotes raw scores for a specific cell cycle phase along the dividing pseudotime. G0 cells have no dominant phase scores for any phase. Heatmap (lower panel) showing expression levels of individual genes identified as phase-specific by the algorithm. Each column is a cell and each row is a gene. Such ordering is largely consistent with the original cell identities by the clustering analysis as shown in the bottom color-coded bar. (C) Dot plots showing expression levels of phase-specific genes along microglia dividing pseudotime. Genes for each phase are plotted in separate graphs with each dot representing the level of expression for a given gene in a given cell. Curves show average expression of all genes assigned to a phase along dividing pseudotime. Expression of microglial signature genes is shown at the bottom. (D) The gray pie chart showing overlaps of phase-specific genes identified by the algorithm compared with four published cell cycle gene sets. The “Novel” category means the percentage of genes found here that were not reported in any of the four datasets. Colored pie charts are breakdowns of the genes from each category based on the phase assignment. (E) Table showing gene names identified as “Novel” in (D). Genes that may play a role in cell division are in bold. (F) Heatmap (upper panel) showing pseudotime ordering of P7-C0 microglia by normalized phase scores. The bottom panels use *Ankle1* as an example to show its expression dynamics along dividing pseudotime. The smoothened expression is the average expression of *Ankle1* along the cell order computed with a fixed window size (length of ordered cells/10). *Ankle1* peaks at G2/M and M. Each dot denotes its expression level for a single cell. See also Figure S3, Figure S4, Table S4.

Microglia in the first two postnatal weeks are highly proliferative (Nikodemova et al., 2015). Previous work showed that in scRNA-seq analysis, cell cycle genes can overload the major principal components that underlie cell-to-cell variations, thereby masking other, more functionally relevant, differences between cells (Buettner et al., 2015). Indeed, many cell cycle genes ranked at the top of the first 3 principal components explaining variations in P7 microglia, and using conserved cell cycle genes for clustering, these cells were clearly segregated based on their presumable cell cycle phases (Figures S3A and S3C). To eliminate this confounding factor and unmask the underlying microglial heterogeneity, we used an established algorithm to regress out cell cycle effects followed by re-clustering of these P7 microglial cells (Figures S3B and S3C; see the Methods). This approach yielded three clusters: P7-C0, P7-C1 and P7-C2 (Figure 4A). Interestingly, these three clusters largely corresponded to the three dominant cell cycle-independent clusters (cluster 1, 2 and 5) found in the original analysis, and the two original cycling clusters (cluster 3 and 4) were simply re-distributed among P7-C0, P7-C1 and P7-C2 following cell cycle regression (Figure 4A). These findings suggested that, irrespective of cell cycle states, there were only 3 categories of microglia at the P7 stage, and that the two major clusters (P7-C0 and P7-C1) were each comprised of non-dividing cells and their corresponding dividing cells in two cell cycle phases (Figure 4A).

It is of great interest to understand the gene regulation of microglial division because microgliosis is prevalent in disease and injury. We took advantage of the scRNA-seq data that presumably included gene expression information of individual dividing microglia at various phases of the cell cycle, and computationally reconstructed this dividing process (Figures 4B, S4A and S4B; see the Methods). To obtain cell cycle phase-specific gene sets for endogenous microglia, we analyzed the P7 microglia (P7-C0 and P7-C1 were analyzed separately) due to their sufficient population sizes, although we could also detect 11 dividing adult microglia in our data (Figure 1E). We used lists of conserved genes from a well-established dataset for HeLa cell lines as the “seeds” to reconstruct 5 cell cycle phases along microglial division pseudotime (see also Methods) (Macosko et al., 2015; Whitfield et al., 2002). Then the algorithm searched for other genes correlated with phases and refined the ordering of cells based on updated phase-specific gene expression. Through a number of repetitions, lists of genes stably assigned to each phase were obtained (Figures 4B, S4B, and S4D; Table S4). Averaged expression of these gene sets showed a “wave-like” pattern along the dividing pseudotime, whereas microglial signature genes were expressed at a consistent level (Figures 4C and S4C).

In total, we identified 315 periodically expressed genes for P7-C0 cells, and 347 genes for P7-C1 cells (some genes were expressed in more than one phase) (Table S4). Excluding the seeds input, 254 (81% of 315) genes for P7-C0 and 227 (65% of 347) genes for P7-C1 were identified by the algorithm, and 183 genes overlapped in both clusters. We compared these machine-reported gene lists with four published datasets for experimentally characterized cell cycle genes (including the HeLa data we used as seeds), and found that the majority of newly identified genes appeared in at least one of the four datasets (Figures 4D and S4E), suggesting the robustness of this analysis. More importantly, we identified many genes that have yet to be described as periodically expressed in dividing cells, although more than half of these new genes have possible links to cell proliferation based on gene function annotations (Figures 4E and S4F). For example, we discovered genes that are involved in DNA damage response and repair (*Ankle1, Lig1, Rpa1*), histone mRNA decay (*Eri1*), and genes with epigenetic functions such as replication-independent histones (*H2afz, H3f3b*) and chromatin modifiers (*Hmgn2, Dnmt1*) (Figure 4F).

### Heterogeneity of postnatal microglia revealed by scRNA-seq

To understand early postnatal microglial heterogeneity, we next examined three P7 clusters (after cell cycle regression). Interestingly, P7-C2 was a small cluster that mainly included cells clustered together with embryonic E14.5 microglia (cluster 5) in the initial analysis, suggesting the presence of primitive microglia in the postnatal brain (Figures 1C and 5A). Cells belonging to P7-C2 expressed microglial signature genes at lower levels (e.g. *Tmem119, Selplg, P2ry12, Tgfbr1*), and displayed up-regulation of genes in metal homeostasis (*Fth1, Ftl1, Mt1*), actin cytoskeleton dynamics (*Tmsb4x, Pfn1, Cfl1*), and ribosomal components (e.g. *Rps14, Rps18, Rpl35, Rps29*) (Figure 5B; Table S5).

**Figure 5.**
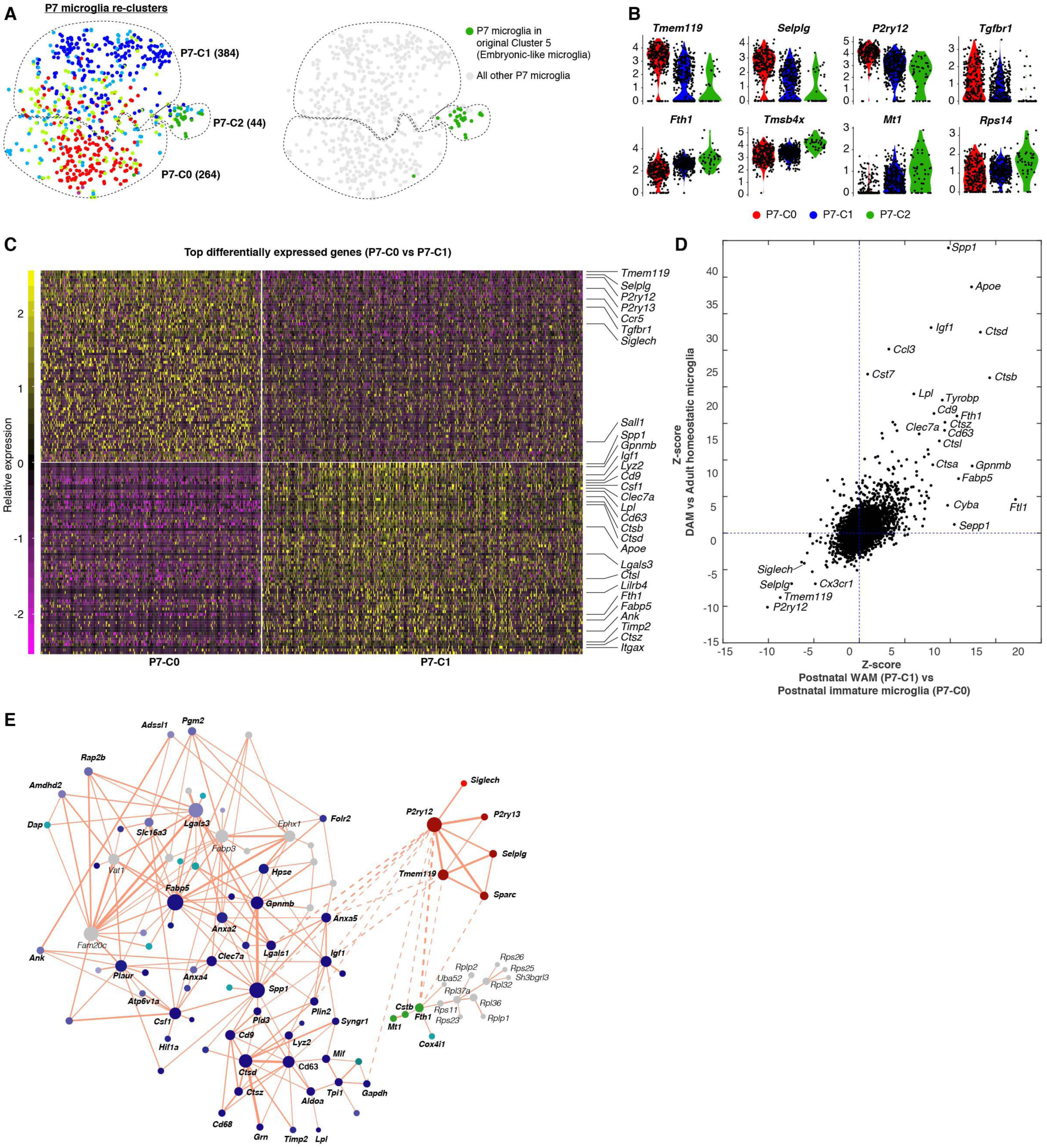
Heterogeneity of postnatal microglia revealed by scRNA-seq. (A) tSNE plot (left panel) showing re-clustering result of P7 microglia as delineated in Figure 4A (color coding is the same). Numbers of cells in each cluster are given in parentheses. tSNE plot on the right highlight P7 cells in the original embryonic-like cluster (cluster 5), showing that P7-C2 in the re-clustering are embryonic-like cells in the P7 brain. (B) Violin plots showing some differentially expressed genes between P7-C2 and the other two clusters (down-regulated genes are shown at the top; up-regulated genes are shown at the bottom). (C) Heatmap showing top 70 differentially expressed genes between P7-C0 and P7-C1. P7-C0 have higher expression levels for homeostatic genes and P7-C1 enrich many disease-associated genes. Genes that have been documented as mis-regulated in disease settings are labeled on the right. (D) Comparison of gene expression changes in DAM (relative to homeostatic microglia) with the changes in P7-C1 (relative to P7-C0) showing similar sets of up-and down-regulated genes in two cases. Genes that have been associated with diseases are labeled. (E) Gene network showing correlated gene modules underlying cluster identities of P7 microglia (See also Methods). Each gene is colored based on its differential expression levels among three P7 clusters (i.e. marker genes for P7-C0 are in red; marker genes for P7-C1 are in blue; marker genes for P7-C2 are in green; genes that are up-regulated in both P7-C1 and P7-C2 are in turquoise; genes that are not differentially expressed are in gray). Colors of different shades and tints indicate the significance levels of the changes. The size of a circle represents how well connected a gene is with other genes. The thickness of an edge denotes the level of correlation between two genes. Solid lines stand for positive correlation and dashed lines stand for negative correlation. See also Table S5.

The majority of P7 microglia fell into either P7-C0 or P7-C1 cluster. Microglia from both clusters expressed homeostatic genes, such as *Tmem119, P2ry12, Tgfbr1, Siglech*, and *Sall1*, but P7-C1 cells expressed them at lower levels (Figure 5C; Table S5). Remarkably, P7-C1 microglia exhibited characteristic expression of many genes that have recently been shown to be up-regulated in degenerative disease-associated microglia (DAM) (Keren-Shaul et al., 2017; Krasemann et al., 2017). These included *Spp1, Gpnmb, Igf1, Clec7a, Lpl, Cd9, Cd63, Lgals3, Fabp5, Itgax, Apoe* and *Tyrobp* (Figure 5C). When we directly compared differentially expressed genes in P7-C1 (relative to P7-C0) with those mis-regulated in DAM, we found that they were strikingly similar in that up-regulation of the disease genes was accompanied by down-regulation of the homeostatic gene cassette (Figure 5D). To obtain a better understanding of the genes underlying P7 microglial heterogeneity, we generated a gene network based on correlation between pairs of genes in all P7 microglia (Figure 5E; see the Methods). We highlighted the nodes with colors corresponding to the significance levels in differential gene expression analysis and cluster signatures they represented. We found that three interconnected gene modules explained P7 microglia identities and, more importantly, many DAM signature genes occupied the “hub” positions in the network.

### Validation of postnatal developing white matter-associated microglia

We were fascinated by the resemblance of the P7-C1 microglia to DAM in gene expression, and examined these postnatal microglia in greater detail. We noticed that the top ranked marker genes, *Spp1* and *Gpnmb*, also distinguished this microglial subset from all other myeloid cell populations (Figures 6A and S5A; Table S5). Surprisingly, RNA *in situ* labeling demonstrated that expression of these two genes overlapped almost exclusively in the corpus callosum and cerebellar white matter, and that *Spp1*^*+*^*Gpnmb*^*+*^ cells were positive for the microglia marker *Cx3cr1* (Figure 6B). scRNA-seq data suggested that this P7-C1 population also had higher levels of *Igf1* and *Itgax* expression. Indeed, we detected co-expression of both transcripts in *Gpnmb*-positive cells (Figures 6C and S5B). Because the *Gpnmb*-labeled P7-C1 microglia were intermingled with *Mbp*^+^ oligodendrocytes in the developing white matter (Figure 6D), we named them postnatal white matter-associated microglia (WAM).

**Figure 6.**
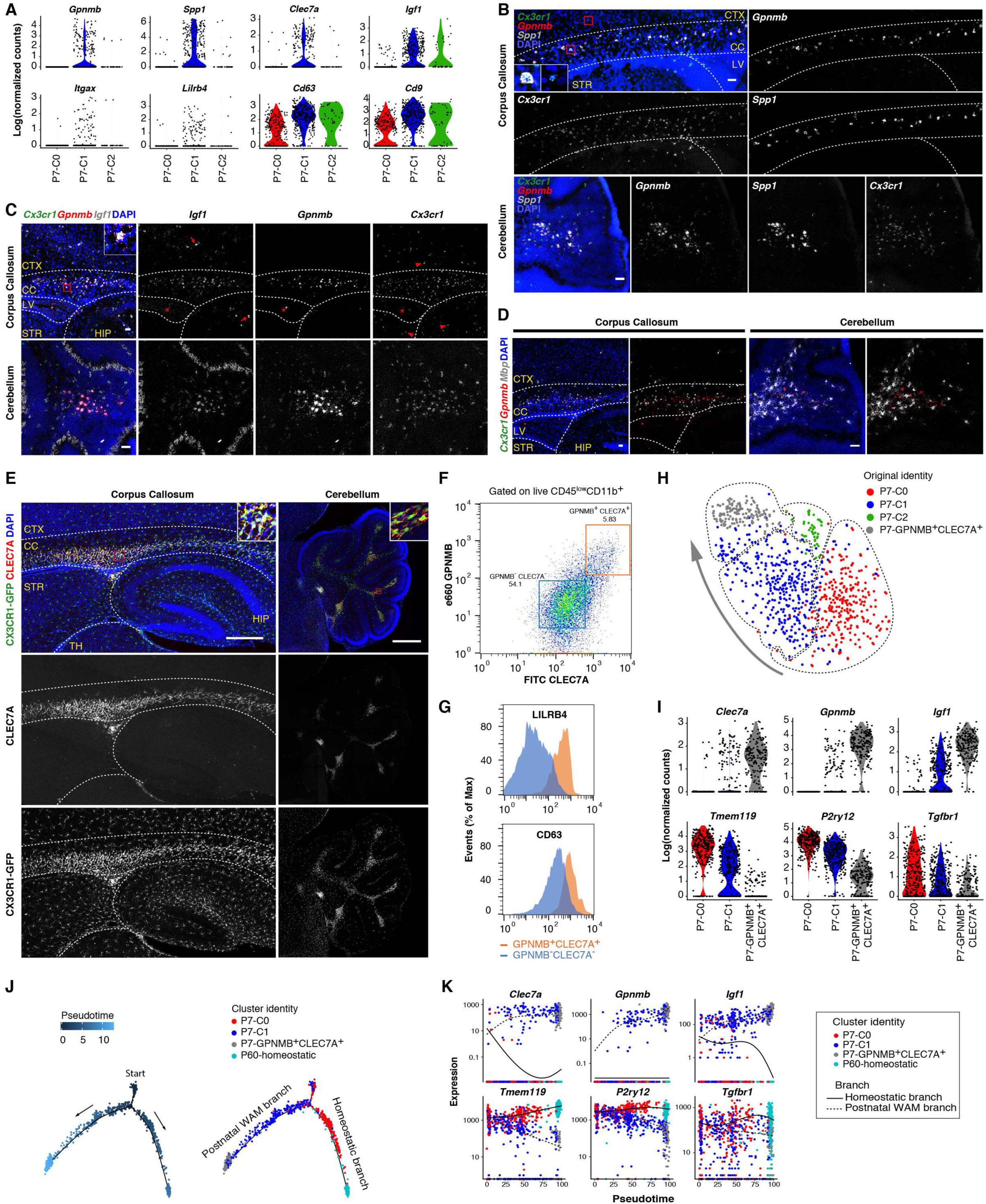
Validation of postnatal developing white matter-associated microglia. (A) Violin plots showing some top up-regulated genes in P7-C1 compared with the other two clusters. All of these genes have been linked to degenerative disease-associated microglia. (B) RNA *in situ* (RNAscope) validation of *Spp1* and *Gpnmb* expression in P7-C1 microglia. *Spp1* and *Gpnmb* signals overlap in the corpus callosum and white matter region of cerebellum, and these cells are also positive for the microglia marker *Cx3cr1*. RNA *in situ* (RNAscope) validation of *Igf1* expression in P7-C1 microglia. *Gpnmb*^*+*^*Cx3cr1*^*+*^ microglia are positive for *Igf1* in the corpus callosum and white matter region of cerebellum, and *Gpnmb*^*+*^*Cx3cr1*^*+*^*Igf1*^*+*^ cells are also seen in the ventricles (asterisk). Microglia in the cortex, hippocampus and striatum are negative for *Gpnmb* and *Igf1* (arrow heads). *Igf1*^*+*^*Cx3cr1*^*-*^neural cells are also present in the cortex and hippocampus (arrows). (D) RNA *in situ* (RNAscope) showing *Gpnmb*^*+*^*Cx3cr1*^*+*^ microglia are intermingled with *Mbp*^*+*^ oligodendrocytes in the developing white matter. (E) Immunohistochemistry validation of CLEC7A expression by postnatal white matter-associated microglia (WAM). CLEC7A^+^CX3CR1-GFP^+^ microglia are populated in the developing white matter and near ventricles. (F) FACS plot showing P7 cerebellar GPNMB^+^CLEC7A^+^ cells (gated on live CD45^low^CD11b^+^ single cells; orange box) isolated for scRNA-seq. Double negative cells (blue box) are used for comparisons in (G) by flow cytometry. (G) Histograms showing higher levels of LILRB4 and CD63 surface expression in GPNMB^+^CLEC7A^+^ microglia compared with GPNMB-CLEC7A-cells. (H) tSNE plot showing clustering result (dashed circles) after combining GPNMB^+^CLEC7A^+^ cells with the originally sequenced P7 microglia. Cells are color coded based on the identities of three previously identified P7 clusters and the newly isolated double positive cells. The arrow indicates gradual changes of transcriptomes from postnatal immature state (P7-C0) towards more polarized GPNMB^+^CLEC7A^+^ state. (I) Violin plots showing further up-or down-regulation of differentially expressed genes in P7-GPNMB^+^CLEC7A^+^ compared with P7-C1 microglia by scRNA-seq. Notably, almost 100% of the double positive cells expressed the *Igf1* marker. (J) Pseudotime analysis of P7 microglia together with P7-GPNMB^+^CLEC7A^+^ and P60 homeostatic microglia showing developmental trajectories from P7-C0/P7-C1 mixed starting point to P7-GPNMB^+^CLEC7A^+^ postnatal WAM branch (via P7-C1) and to P60 homeostatic branch (via P7-C0). Each dot is a cell. (K) Gene expression dynamics for two trajectories in (J) along developmental pseudotime, when the postnatal WAM branch gradually turns on disease-associated genes and down-regulates homeostatic genes. The homeostatic branch displays the opposite trends of gene expression. Scale bars: 50um in (B)-(D), 500um in (E). CTX: cortex; CC: corpus callosum; LV: lateral ventricle; STR: striatum; HIP: hippocampus; TH: thalamus. See also Figure S5, Table S5.

Intriguingly, we discovered that postnatal WAM and periventricular myeloid cells specifically expressed CLEC7A protein (Figure 6A, 6E, and S5A), which labels microglia surrounding amyloid plaques in mouse models of Alzheimer’s disease (AD) and has been described as a marker for DAM (Keren-Shaul et al., 2017; Krasemann et al., 2017). To further characterize postnatal WAM, we stained P7 microglia with the surface proteins, CLEC7A and GPNMB, and sorted the double positive cells by FACS (Figure 6F). GPNMB^+^CLEC7A^+^ cells also expressed higher levels of LILRB4 and CD63 which were part of the P7-C1 RNA signature and further established the identity of this microglial population (Figures 6A, 6F, and 6G). Additionally, scRNA-seq of microglia freshly isolated based on GPNMB and CLEC7A co-expression validated the original gene expression pattern observed with the initial P7 clustering results; the clustering algorithm also reported another GPNMB^+^CLEC7A^+^ concentrated cluster which overlapped with a portion of P7-C1 cells (Figure 6H and 6I). This new double positive cluster expressed all the markers identified in P7-C1, just at higher levels on average and in a much higher percentage of the cells (Figure 6I; Table S5). This was possibly due to the enrichment of this postnatal WAM subset at their most polarized state. Consistent with this interpretation, *Spp1* was detected in close to 100% of the double positive cells by scRNA-seq, and SPP1 only labeled a portion of CLEC7A^+^ cells on tissue sections, presumably representing the most typical form of postnatal WAM (Figure S5C). Interestingly, postnatal WAM also displayed distinct cytokine/chemokine secretion profiles, including *Ccl3* and *Ccl4* which have been linked to microglia in aging and AD models (Kang et al., 2018) (Figure S5D).

Because there seemed to be a gradient shift in overall gene expression from P7-C0, to P7-C1, to P7-GPNMB^+^CLEC7A^+^ cells (Figure 6H), we performed developmental pseudotime analysis to computationally reconstruct possible developmental trajectories of postnatal microglia. We discovered that one trajectory pointed towards the development of postnatal WAM, while the other pointed towards homeostatic, mature cells (Figure 6J). These data suggest that postnatal immature microglia can either turn on genes specific for classical microglia found in the adult brain (P60-homeostatic) or assume the WAM phenotype (P7-C1 and P7-GPNMB^+^CLEC7A^+^) similar to DAM in aging and neurodegeneration (Figure 6K).

### Transient appearance of WAM with peak at P7, independent of TREM2 or APOE

To investigate the temporal and spatial appearance of postnatal WAM during development, we stained brain sections from E17.5 to P60 with CLEC7A antibodies. In general, we discovered that CLEC7A^+^ microglia were associated transiently with the developing white matter, and found in regions of neurogenesis. During the late embryonic (E17.5) stage, CLEC7A^+^ microglia appeared almost exclusively in the neurogenic niches near the lateral ventricles (Figure 7A), while CLEC7A-IBA1^+^ cells were present throughout the brain parenchyma. Starting at P4, CLEC7A^+^ microglia populated the corpus callosum and cerebellar white matter (Figure 7B), and they continued to expand and peaked around P7 (Figures 6E and 7E). By P14, these white matter-associated microglia were barely detectable (Figures 7C and 7E). Interestingly, starting at P14 and becoming more prominent during adulthood, CLEC7A^+^ microglia were also observed in the hippocampal dentate gyrus region, the subventricular zone and along the rostral migratory stream where adult neurogenesis and precursor migration occur (Figures 7C and 7D). The presence of CLEC7A^+^ microglia in these strategic niches indicated that they might be involved in embryonic and adult neurogenesis. Morphologically, CLEC7A^+^ microglia in the P7 white matter were amoeboid in shape and had thicker primary branches with larger cell bodies (Figures 7F-7H). By contrast, CLEC7A-microglia in the cortex were much more ramified. These CLEC7A^+^ cells also congregated at a higher density compared with CLEC7A^-^ immature microglia (Figures 7F, 7H, and S6A). Taken together, these data suggest that postnatal WAM represent a specialized population which transiently appears in the developing white matter, rather than a general immature state for all microglia.

**Figure 7.**
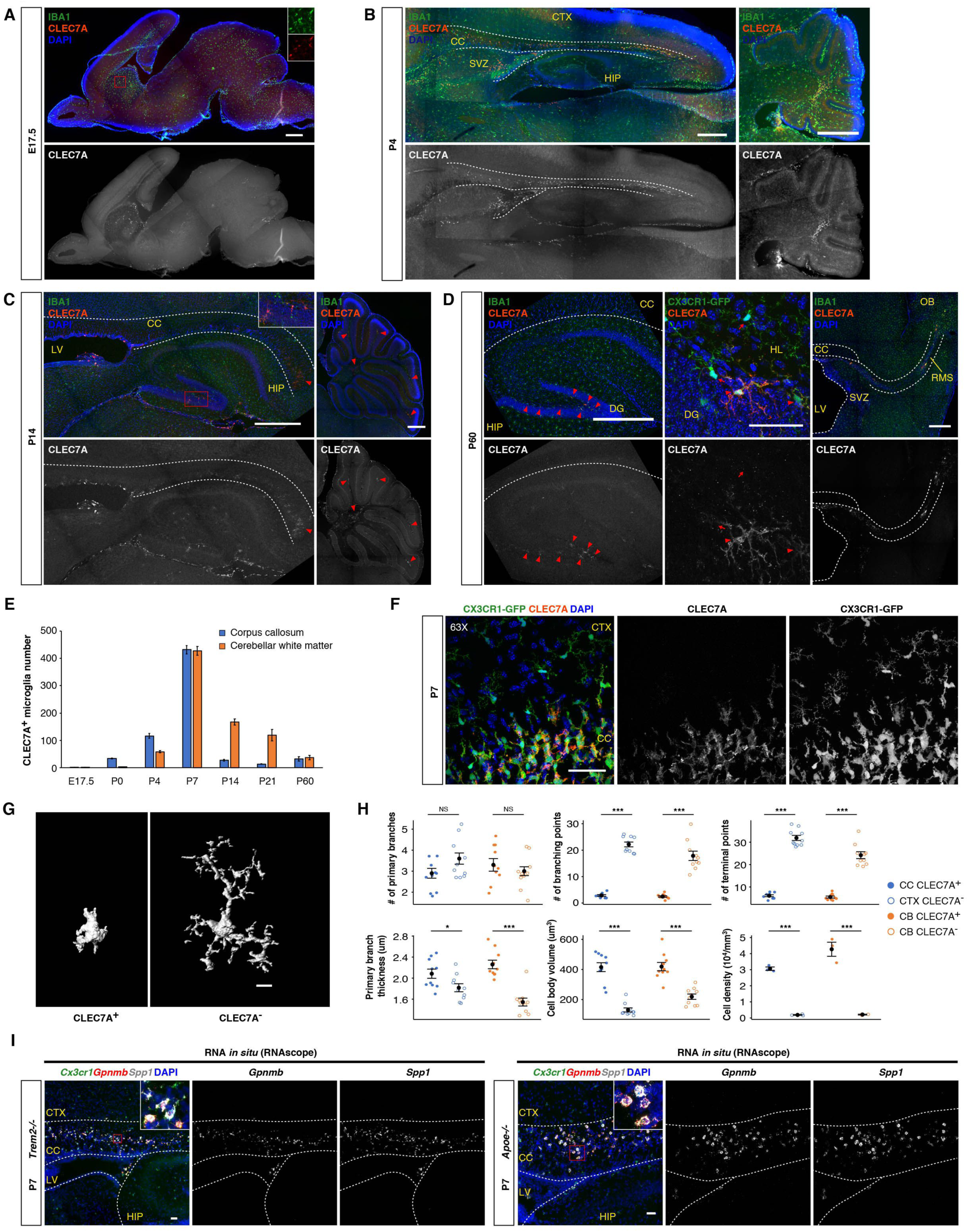
Transient appearance of amoeboid postnatal WAM, independent of TREM2-APOE regulation. (A)-(D) Immunohistochemistry on CLEC7A expression from different developmental stages. (A) E17.5 brain sections showing lack of CLEC7A^+^ cells in the corpus callosum and cerebellum, and their presence in the subventricular region (inset). (B) CLEC7A^+^ microglia start to populate in developing white matter at P4. They are also seen in the subventricular region. (C) At P14, only a few microglia with weak CLEC7A signals are seen in the white matter (arrow heads). Inset shows CLEC7A^+^ microglia in hippocampal dentate gyrus. (D) At P60, CLEC7A^+^ microglia are mostly absent in the white matter, but show in hippocampal dentate gyrus (arrow heads) and neurogenic regions (SVZ and RMS). Middle panels: 63X confocal images showing CLEC7A^+^ (arrow heads) and CLEC7A-(arrows) microglia in the hippocampus. (E) Quantification of CLEC7A^+^ microglia on 50um sagittal brain sections across developmental stages. n=3 for each stage. (F) 63X confocal images showing differences in morphology and density between CLEC7A^+^ and CLEC7A-microglia at P7. (G) 3D reconstruction of representative P7 microglia showing amoeboid morphology of CLEC7A^+^ microglia. (H) Quantification showing differences in morphology (n=10 each) and density (n=3 each) between CLEC7A^+^ and CLEC7A-microglia at P7. *** *P*<0.001, ** *P*<0.01, * *P*<0.05. One-way ANOVA followed by pairwise t-test with Bonferroni correction. (I) RNA *in situ* (RNAscope) showing presence of *Gpnmb*^*+*^*Spp1*^*+*^ microglia in the corpus callosum of *Trem2* or *Apoe* knockout mice at P7. CTX: cortex; CC: corpus callosum; SVZ: subventricular zone; LV: lateral ventricle; HIP: hippocampus; DG: dentate gyrus; HL: hilus; RMS: rostral migratory stream; OB: olfactory bulb. Scale bars: 500um in (A)-(D), except for 50um in the middle panels of (D); 50um in (F) and (I); 10um in (G). Data are represented as mean ± SEM in (E) and (H). See also Figure S6.

It has been shown that transcriptional changes in degenerative disease-associated microglia depend on the TREM2-APOE pathway, and the DAM signature genes such as *Spp1, Gpnmb* and *Clec7a* are suppressed in disease models that also lack *Trem2* or *Apoe* (Keren-Shaul et al., 2017; Krasemann et al., 2017). Surprisingly, despite their similarity in gene expression to DAM, amoeboid *Gpnmb*^*+*^*Spp1*^*+*^CLEC7A^+^ microglia were still densely populated in the P7 white matter of *Trem2* or *Apoe* knockout mice (Figures 7I and S6B-6E). These data suggest that, unlike DAM, postnatal WAM do not depend on TREM2 or APOE for their polarization to occur.

### Phagocytosis of newly formed oligodendrocytes by postnatal WAM

To investigate the biological functions of postnatal WAM, we followed up on our histological observation that CLEC7A^+^ microglia in the developing white matter had amoeboid shapes and frequently contained pyknotic nuclei (Figures 8A and 8B; Movie S1). Amoeboid morphology has indeed been associated with a phagocytic state in microglia or macrophages (Bohlen et al., 2017; Ling and Wong, 1993). To compare the phagocytic capability between CLEC7A^+^ and CLEC7A-microglia in the postnatal brain, we exposed acutely sectioned brain slices to pH-sensitive beads and quantified percentages of microglia that engulfed fluorescent beads based on CLEC7A expressivity and brain regions. We found that only CLEC7A^+^ microglia in the corpus callosum and cerebellum preferentially phagocytosed beads (Figures 8C, 8D, S7A, and S7B), and this phenomenon was independent of opsonizing factors or other components in serum (Figure 8D). These results suggest that postnatal WAM are highly phagocytic.

**Figure 8.**
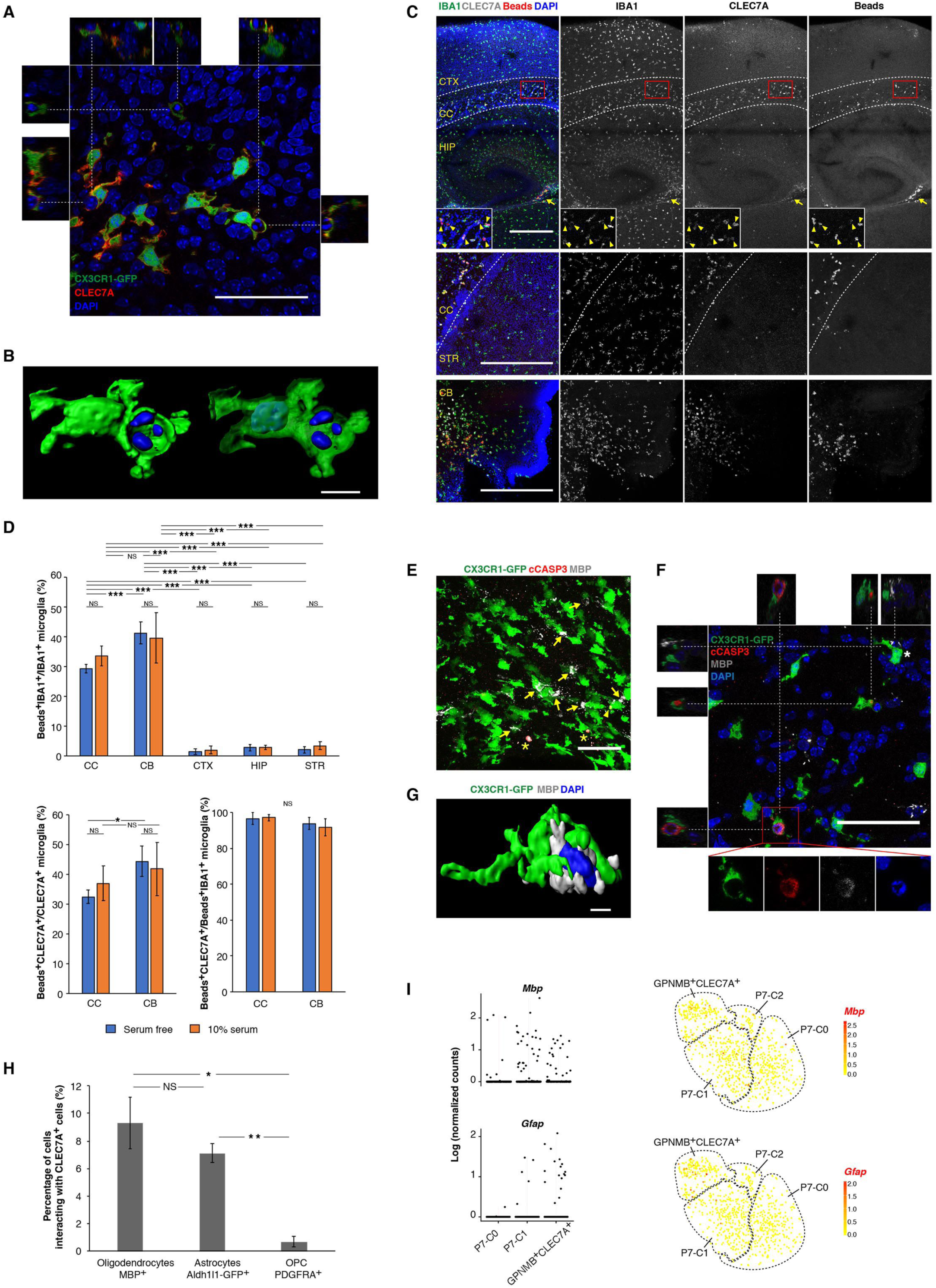
Phagocytosis of newly formed oligodendrocytes by postnatal WAM. (A) Single optical section of a confocal image showing engulfment of nuclei (some with fragmented DAPI signals) by CLEC7A^+^ microglia in the P7 cerebellar white matter. Images from the X-Z and Y-Z axes are shown on the top and sides, respectively. (B) 3D reconstruction of a CLEC7A^+^ microglia engulfing a pyknotic nucleus. The microglia nucleus can be seen by transparent rendering for the CX3CR1-GFP (green) signal on the right. (C) Immunohistochemistry on brain sections after 4hr *in vitro* culturing with pHrodo Zymosan beads. Many CLEC7A^+^ microglia that phagocytosed beads are seen in the corpus callosum (arrow heads) and cerebellar white matter but they are rarely seen in other regions. CLEC7A^+^ cells near ventricles (arrows) also phagocytosed beads. (D) Quantification of the phagocytosis assay in (C). n=5 sections (3 fields for each section). *** *P*<0.001, ** *P*<0.01, * *P*<0.05. Two-way ANOVA followed by Tukey’s multiple comparisons test. (E) Immunostaining of P7 corpus callosum region showing extensive interactions between postnatal WAM and MBP^+^ oligodendrocytes (arrows). Some cleaved Caspase 3 signals overlap with MBP signals (asterisks). cCASP3^+^ inclusion is seen in microglial cells (arrow head). (F) Single optical section of a confocal image showing engulfment of dying (cCASP3^+^) MBP^+^ oligodendrocytes by CLEC7A^+^ microglia in the P7 corpus callosum. Images from the X-Z and Y-Z axes are shown on the top and left, respectively. At the upper right corner, a microglial cell (asterisk) physically contacts a cCASP3-MBP^+^ oligodendrocyte, which is 3D reconstructed in (G). (H) Quantification for percentage of each cell type that interacts with CLEC7A^+^ cells in the corpus callosum. Interaction is defined as >30% of cellular volume overlapping with the CLEC7A (or CX3CR1-GFP) signal. n=3 sections each. ** *P*<0.01, * *P*<0.05. One-way ANOVA followed by pairwise t-test with Bonferroni correction. (I) Detection of *Mbp* and *Gfap* transcripts in postnatal WAM by scRNA-seq. These two gene transcripts are disproportionally found in P7-C1 and particularly P7-GPNMB^+^CLEC7A^+^ microglia. tSNE plots (same as in Figure 6H) on the right highlight cells that have detectable *Mbp* or *Gfap* expression. CTX: cortex; CC: corpus callosum; HIP: hippocampus; STR: striatum; CB: cerebellum. Scale bars: 50um in (A), (E) and (F); 500um in (C); 10um in (B) and (G). Data are represented as mean ± SEM in (D) and (H). See also Figure S7, Table S6, Table S7, Movie S1, Movie S2.

Since the occurrence of postnatal WAM effectively correlates with the onset of CNS myelination when newly formed oligodendrocytes undergo massive cell death (Barres et al., 1992; Trapp et al., 1997), we hypothesized that postnatal WAM might phagocytose some of these oligodendrocytes. Labeling oligodendrocytes with MBP in the CX3CR1-GFP mouse showed that microglia extensively interacted with star-shaped MBP^+^ cells in the developing white matter (Figure 8E). More importantly, we observed postnatal WAM containing cleaved Caspase-3 (cCASP3)-positive inclusions, which were MBP positive and possessed pyknotic nuclei (Figures 8E, 8F, and S7E; Movie S2). These data suggest that postnatal WAM phagocytose dying newly formed oligodendrocytes during myelination. Interestingly, cCASP3-negative oligodendrocytes were frequently “hugged” by postnatal WAM, raising the possibility of them actively contributing to elimination of cells (Figure 8G). Moreover, we rarely observed juxtaposition of PDGFRA^+^ oligodendrocyte precursor cells (OPCs) with postnatal WAM (Figures 8H and S7C), suggesting that this phagocytic activity mainly targets newly formed oligodendrocytes but not their precursors. In addition, we could not find cCASP3^+^ Aldh1l1-GFP labeled astrocytes, although substantial interactions between astrocytes and postnatal WAM existed (Figures 8H, S7D, and S7F). Remarkably, *Mbp* and *Gfap* transcripts were detected in certain percentages of P7-C1 (32/323=9.9% for *Mbp*; 5/323=1.5% for *Gfap*) and P7-GPNMB^+^CLEC7A^+^ microglia (26/192=13.5% for *Mbp*; 17/192=8.9% for *Gfap*) by scRNA-seq, while *Pdgfra* counts were zero in all P7 microglia (Figure 8I). Because the detection rates for P7-C0 microglia (8/274=2.9% for *Mbp*; 2/274=0.7% for *Gfap*) were more than 2 folds lower for both genes (*P*<0.001 for either gene, Chi-squared test), these differences were unlikely due to mRNA contamination during cell isolation. The simplest explanation is that scRNA-seq sufficiently captured transcripts from engulfed oligodendrocytes (*Mbp*) or astrocytes (*Gfap*) in postnatal WAM. Taken together, these data suggest that postnatal WAM phagocytose newly formed oligodendrocytes and perhaps to a lesser extent, astrocytes.

## Discussion

Microglia and related brain myeloid cells are now recognized as functional modulators of developmental processes and neurodegeneration (Li and Barres, 2017). It is commonly accepted that heterogeneous myeloid populations are involved throughout the course of disease progression (Prinz et al., 2011). The extent of microglial heterogeneity and its functional significance in the mature homeostatic brain, however, remains elusive. The picture is even less clear when the additional variable of developmental stages is considered. In this study, we used deep single-cell RNA sequencing (scRNA-seq) to address the question of microglial heterogeneity in the broad context of innate immune cells, and we systematically analyzed transcriptomes of these cells from distinct brain regions and across developmental time points.

We utilized a semi-automated Smart-seq2 platform to perform deep sequencing for individual cells. The superb sensitivity and accuracy of this scRNA-seq technology compensates for its limitation in throughput (Svensson et al., 2017; Ziegenhain et al., 2017), allowing us to robustly identify cell populations with fewer than 20 cells. Importantly, this approach provides a better chance for detecting low abundance genes such as transcription factors. With its unprecedented depth, this resource will be useful to generate hypotheses for cellular origins of certain biological or pathological processes, which in turn could be tested with the molecular markers identified here for cell purification or functional manipulation.

Our scRNA-seq data along with histology validation showed that adult and early postnatal choroid plexus macrophages (CP MΦ) were distinctively different in their gene expression profiles. For example, we found that high MHC-II expression, previously associated with CP MΦ (Prinz et al., 2017), was only detected in the adult brain. Interestingly, it has been shown that immune response in the choroid plexus is also altered in the aged brain (Baruch et al., 2014), suggesting dynamic changes of CP cells across lifespan. It will be interesting to identify the signals that contribute to this phenotypic change as well as its functional relevance.

Genes upregulated in proliferative cells are highly correlated with cell cycle genes in tumor and many other cell types (Whitfield et al., 2002). *In vivo* studies of periodical gene expression through cell cycle stages have been challenging due to the limited numbers of cycling cells that can be obtained and lack of phase-specific markers. We took advantage of scRNA-seq data as a random sampling of individual cells along a continuum, and reconstructed molecular features throughout microglial cell division without the need of artificial synchronization. We generated cell cycle phase-specific gene sets for microglia which will not only provide a fresh perspective on our understanding of microglial proliferation, but also serve as a valuable resource for studies of mammalian cell cycle.

Using this deep sequencing approach, we discovered surprisingly little transcriptomic heterogeneity among adult homeostatic microglia. The one cluster distinctively separated from other homeostatic cells showed elevated expression of immediate early genes, which we attributed to isolation artifacts. Our data point to a lack of prominent transcriptomic differences among classical adult microglia even across varying brain regions, a finding we confirmed by bulk RNA-seq of highly purified region-specific homeostatic (TMEM119^+^) microglia. These results indicate that the previously observed regional differences in adult microglia might mainly arise from genes expressed by small percentages of TMEM119^low/-^microglia (e.g. residual CLEC7A^+^ cells shown in Figure 7C-7E), which were gated out in our bulk RNA-seq analysis (Ayata et al., 2018). Consistent with this interpretation, many genes (*Apoe, Igf1, Lilrb4, Lyz2, Colec12, Msr1, Map1lc3b*) that were shown to be part of the phagocytic signature in adult cerebellar microglia (Ayata et al., 2018) were also up-regulated in postnatal WAM (Table S5). Meanwhile, other genes previously attributed to microglial regional heterogeneity could be due to differential abundance of non-microglial populations in bulk samples with the CD11b selection (Grabert et al., 2016). Furthermore, additional brain regions may uncover transcriptomic differences not present in the 4 major regions tested here. For example, microglia in the midbrain have been shown to possess distinct gene expression profiles compared with cortical microglia (De Biase et al., 2017). Microglial heterogeneity can also be represented by measurements other than gene transcription, such as electrophysiological properties, epigenetic features or functional readouts (Ayata et al., 2018; De Biase et al., 2017).

In sharp contrast to microglia in the adult stage, early postnatal microglia are much more heterogeneous. First, we captured proliferative microglia with dividing cell signatures as discussed. Second, we detected a small cluster of P7 cells that resembled E14.5 microglia. Whether this population merely reflects developmental asynchrony or holds a functionally important subset with certain stemness features awaits future investigation. Third, we identified a subset of microglia that we named postnatal white matter-associated microglia (WAM). We validated the presence of postnatal WAM at both RNA and protein levels using different methods, and identified *Gpnmb, Spp1* and *Clec7a* expression as characteristic molecular markers for these cells. We also demonstrated that GPNMB and CLEC7A surface antigens could be used to highly purify postnatal WAM, as nearly 100% of these sorted cells expressed the WAM gene signature. Comprehensive analyses of these cells in comparison with the other dominant population of immature microglia showed that postnatal WAM were amoeboid, accumulated at a higher density, and displayed gene expression patterns that mimic degenerative disease-associated microglia (DAM) (Keren-Shaul et al., 2017; Krasemann et al., 2017).

The parallels between DAM and postnatal WAM are particularly interesting as they support the concept that genes expressed during development are reactivated in aging and degeneration (Hong et al., 2016). Functional characterization of postnatal WAM may thus provide valuable clues for understanding pathophysiology of neurodegenerative diseases. Indeed, while postnatal WAM engulf newly formed oligodendrocytes and possibly newborn astrocytes during development, DAM are also phagocytic and involved in phagocytosis of Aβ plaques in AD models (Keren-Shaul et al., 2017; Krasemann et al., 2017). In addition, postnatal WAM enrich many metabolic genes including almost the entire molecular machineries for oxidative phosphorylation, glycolysis and beta oxidation (Figure S7G; Tables S6 and S7). We also found up-regulation of a large cohort of genes responsible for lysosomal acidification, lipid transport and metabolism, as well as lysosomal enzymatic functions (Table S7). This indicates high levels of energy turnover, particularly lipid metabolism, in postnatal WAM that may be required for phagocytosis of lipid-rich oligodendrocytes. Interestingly, mis-regulated lipid metabolism has also been observed in DAM and in many other disease situations (Keren-Shaul et al., 2017; Yadav and Tiwari, 2014), leading to an open question of how altered metabolic states are linked to disease pathology.

Remarkably, postnatal WAM do not depend on TREM2-APOE signaling to initiate the DAM gene signature. Understanding the molecular underpinnings of postnatal WAM polarization may provide handles to shift microglia in disease settings towards beneficial ones. The presence of DAM-like microglia early in life also raises the tantalizing hypothesis that AD pathogenesis may include a pathogenic re-appropriation of latent microglial functions, triggered by different upstream stimuli.

We showed here that postnatal WAM are not a general intermediate step for microglia development, but specifically and transiently appear in the developing white matter. The timing of their appearance coincides with the onset of myelination, during which large numbers of newly formed oligodendrocytes die (Barres et al., 1992; Trapp et al., 1997). Overproduction of cells, balanced by controlled apoptosis, is a common theme during development of multicellular organisms, although the biological significance behind this is often unknown. Dying or dead cells need to be cleared away by professional phagocytes in order to maintain tissue homeostasis. Alternatively, these phagocytes may promote cell death to sculpt tissue structures and create functional circuits (Fuchs and Steller, 2011). It is thus possible that efficient engulfment of newly formed oligodendrocytes by postnatal WAM in the white matter is necessary to generate evenly spaced oligodendrocyte tiling and to make room for myelination. Besides the phagocytic function we presented here, postnatal WAM may play other roles by interacting with neural or immune cells through secreted molecules. We showed that postnatal WAM display unique cytokine/chemokine profiles and express many trophic factors including *Igf1, Spp1, Lgals1*, and *Lgals3* (Figure S5D). Interestingly, and possibly related to our finding, a CD11c (ITGAX)-expressing microglial subset in the postnatal brain has been suggested to promote CNS myelination via secreted IGF1, which may exert its effect by supporting OPC survival (Hagemeyer et al., 2017; Wlodarczyk et al., 2017).

In summary, profiling individual mouse microglia during late embryonic and postnatal development provides a resource which complements current gene expression datasets of microglia from adult and diseased brains. The extraordinary sequencing depth detected many additional genes expressed in microglia at different developmental stages and brain regions, and led to the discovery of a unique microglial subset termed postnatal WAM. Future functional studies of the identified microglial and myeloid populations will likely provide insights into the pivotal roles innate immune cells play in brain development, homeostasis and disease.

## Acknowledgments

We thank Ryosuke Kita for his effort to make the Myeloid Single-cell RNA-Seq webpage. We thank current and past Barres lab members, particularly Steven Sloan, Christopher Bohlen, Frederick Christian Bennett and Mengmeng Fu for their help and insightful feedback on the manuscript. We thank members of the Wyss-Coray lab, especially Tal Iram and John Pluvinage for their technical support and all others for helpful discussions. We thank David Simon, Jianjin Shi, Wan-Jin Lu, Charles Kwok Fai Chan for discussions and generous help. We owe a debt of gratitude to the Stanford Shared FACS Facility, particularly Brandon Carter and Meredith Weglarz, for their excellent service. We appreciate John Mulholland and Kitty Lee from the Cell Sciences Imaging Facility at Stanford for help with confocal imaging. We are thankful for the assistance of Chi-Li Chiu from BITPLANE with Imaris software. This work is funded by the JPB Foundation (B.A.B.), Adelson Medical Research Foundation (B.A.B.), Vincent J. Coates Foundation (B.A.B.), National Institutes of Health (R01 DA015043, B.A.B.), the Veterans Administration (T.W.-C.), the NOMIS Foundation (T.W.-C), National Institutes of Health (R01 MH110504, G.Y.), the Helen Hay Whitney Foundation (L.O.S.), National Multiple Sclerosis Society (L.O.S.), the Life Science Research Foundation (L.E.C.). Q.L. is supported by a Stanford Medicine Dean’s Fellowship.

## Author Contributions

Conceptualization, Q.L. and B.A.B.; Methodology, Q.L., Z.C., S.D., G.Y.; Software, Q.L., Z.C., G.Y.; Investigation, Q.L., Z.C., L.Z., S.D., N.N.; Formal analysis, Q.L., Z.C., L.Z.; Writing – Original Draft, Q.L., Z.C., L.Z.; Writing – Review & Editing, Q.L., Z.C., L.Z., M.L.B., L.O.S., T.W.-C.; Funding Acquisition, B.A.B.; Resources, S.D., N.N., J.O., S.R.Q., G.G., M.L.B., L.O.S., L.E.C., J.M.; Supervision, B.A.B., G.Y., T.W.-C., S.R.Q.

## Declaration of Interests

The authors declare no competing financial interests.

## Methods

### Animals

All sequencing experiments were performed using C57BL/6 mice of defined ages (Charles River). Trem2 KO mice (C57BL/6J-Trem2em2Adiuj/J) and ApoE KO mice (B6.129P2-Apoetm1Unc/J) were purchased from Jackson Laboratories (027197 and 002052). CX3CR1-GFP mice (B6.129P-*Cx3cr1tm1Litt*/J) were purchased from Jackson Laboratories (005582) and heterozygotes were used for the experiments. Aldh1l1-EGFP mice were used to visualize astrocytes (Chung et al., 2013). Male mice were used in all experiments. Animals were housed and handled in accordance with the guidelines of the Administrative Panel on Laboratory Animal Care of Stanford University.

### Tissue dissection and cell isolation

To isolate cells for scRNA-seq and bulk RNA-seq, P60 mice or pregnant mice with E14.5 embryos were euthanized with CO2, and P7 mice were decapitated directly. For the E14.5 stage, whole brains were dissected out from timed embryos, and collected into 1ml PBS with 1% fetal calf serum (added 10ul Ambion DNase, 2ul RNase inhibitor) on ice. The sample was passed through a 26G needle 5-10 times to create single cell suspension. Cells were spun down (500g) at 4°C for 5min, and suspended in 1ml PBS for FACS staining. For P7 and P60 stages, cortex, cerebellum, hippocampus, striatum, olfactory bulb and choroid plexus were dissected out. For P7 GPNMB/CLEC7A double sorting, only the cerebellum was dissected and used. Tissues from the same region were pooled from multiple mice (3-8 depending on the ages and regions) into a 6cm petri dish with 200ul cold medium A on ice (15mM HEPES, 0.5% glucose in 1 X Hanks’ Balanced Salt Solution (HBSS without phenol red)). Microglia (myeloid cells) extraction was carried out following a published protocol (Bennett et al., 2016). The whole procedure was done on ice with cold buffers. Briefly, tissues were chopped with razor blade into fine pieces and then transferred into a 5ml douncer containing 5ml medium A (added 200ul 12500 units/mL DNase and 10ul recombinant RNase inhibitor). Tissues were dounced with 6-10 full strokes until no visible chunks were present and then filtered into 50ml falcon tubes using 70um strainers to obtain single cell suspension. Cells were washed with medium A and resuspended in 1.8ml MACS buffer with 3.6ul RNase inhibitor (sterile-filtered 0.5% BSA, 2mM EDTA in 1 X PBS). To remove myelin, cells were incubated with 200ul myelin removal beads (MACS Miltenyl Biotec 130-096-433) for 10 min, and loaded onto LD columns (Miltenyi Biotec 130-042-901) assembled on a MACS magnet stand. Cells in the flow through were collected and washed for standard FACS staining.

### Single cell index sorting

To stain the cells for sorting, 5ul mouse Fc block (BD Pharmingen 553142, 1:60) was added into each sample and incubated for 5 min on ice. Then the cells were incubated with antibodies conjugated with fluorophores or primary antibodies for 10 min at room temperature (RT) on shaker. After wash, if necessary, samples were stained with secondary antibodies for 10 min at room temperature and followed by wash. Cells were resuspended in 300ul FACS buffer (sterile-filtered 1%FCS, 2mM EDTA, 25mM HEPES in 1XPBS) with DNase (1:100, Qiagen 79254) and RNase inhibitor (1:500) for single cell index sorting.

Cell sorting/flow cytometry analysis was done on the cell sorter (BD InFlux) at the Stanford Shared FACS Facility. All events were gated with the consecutive gates: (1) forward scatter-area (FSC-A)/side scatter-area (SSC-A) (2) Trigger Pulse Width/ FSC (3) Live-Dead negative (Green for E14.5; Far Red for P7 and P60 (Thermal Fisher Scientific L34960); Propidium Iodide for P7 GPNMB/CLEC7A double sorting) (4) P7/P60: CD45^low^CD11b^+^, CD45^hi^CD11b^+^; E14.5: c-Kit-CD45^+^; P7 GPNMB/CLEC7A: GPNMB^+^CLEC7A^+^(gated on CD45^low^CD11b^+^). Single cells were index sorted into 96-well plates containing 4uL lysis buffer (4U Recombinant RNase Inhibitor, Takara Bio 2313B), 0.05% Triton X-100, 2.5mM dNTP mix (Thermo Fisher Scientific R0192), 2.5uM Oligo-dT30VN (5′- AAGCAGTGGTATCAACGCAGAGTACT30VN-3′). Plates were briefly vortexed, spun down and snap frozen on dry ice. Plates were stored at −80°C freezer until library preparation.

Antibodies used for FACS sorting/flow cytometry: Rat anti-mouse CD117 (c-Kit), APC (clone 2B8) (ThermoFisher Scientific 17-1171-82, 1:200); Rat anti-mouse CD45, PE-Cy7 (clone 30-F11) (Thermo Fisher Scientific 25-0451-82,1:300); Rat anti-mouse CD11b, Brilliant Violet 421 (clone M1/70) (BioLegend 101236, 1:300); Rat anti-mouse F4/80, PerCP/Cy5.5 (clone BM8) (BioLegend 123128,1:100); Rat anti-mouse CD41, PE (BioLegend 133906,1:200); Rabbit anti-mouse TMEM119 (abcam ab210405, 1:400); Goat anti-rabbit Alexa 488 (Thermal Fisher Scientific 11034, 1:300); Rat anti-mouse GPNMB, eFluor 660 (clone CTSREVL) (ThermoFisher Scientific 50-5708-82, 1:20); Rat anti-mouse CD369 (DECTIN-1/CLEC7A) (clone RH1) (BioLegend 144302, 1:50); Mouse anti-rat IgG1 Antibody, FITC (clone MRG1-58) (BioLegend 407406, 1:200); Armenian Hamster anti-mouse CD85k, PE (H1.1) (BioLegend 144904, 1:100); Rat anti-mouse CD63, PE/Cy7 (clone NVG-2) (BioLegend 143910, 1:100).

### Library preparation for scRNA-seq and bulk RNA-seq

For scRNA-seq, sequencing libraries were prepared following the published Smart-seq2 protocol with the aid of liquid handling robotics (Picelli et al., 2014). Briefly, plates with sorted cells were thawed on ice and incubated at 72°C for 3 min in order to anneal RNAs to the Oligo-dT30VN primer. After that, 6ul reverse transcription mixture (95U SMARTScribe™ Reverse Transcriptase (100U/ul, Clontech 639538), 10U RNase inhibitor (40U/ul), 1XFirst-Strand buffer, 5mM DTT, 1M Betaine, 6mM MgCl2, 1uM TSO (Exiqon, Rnase free HPLC purified)) was added into each well, and RT was performed at 42°C for 90 min, followed by 70°C, 5 min. To amplify cDNA, 15ul PCR amplification mix (1X KAPA HIFI Hotstart Master Mix (Kapa Biosciences KK2602), 0.1uM ISPCR Oligo (AAGCAGTGGTATCAACGCAGAGT), 0.56U Lambda Exonuclease (5U/ul, New England BioLabs M0262S)) was added, and the following PCR program was used: (1) 37°C 30 min; (2)95°C 3 min; (3) 21 cycles of 98°C 20 sec, 67°C 15 sec, 72°C 4 min; (4) 72°C 5 min. Amplified cDNA samples were then purified with PCRClean DX beads (0.7:1 ratio, Aline C-1003-50), and resuspended in 20ul EB buffer. cDNA quality was examined with

a Fragment Analyzer (AATI, High Sensitivity NGS Fragment Analysis Kit:1 bp -6000 bp) following the manufacture’s instruction. Samples with sufficient amount of cDNA content (>0.05ng/ul) and normal peaks on the quantification graphs were retained for library preparation. To make libraries, all samples were first diluted down to 0.15ng/ul (only if higher than 0.15ng/ul) in 384-well plates using Mantis Liquid Handler (Formulatrix) and Mosquito X1 (TTP Labtech) with customized scripts. Nextera XT DNA Sample Prep Kit (Illumina FC-131-1096) was used at 1/10 of recommendation volume, with the help of a Mosquito HTS robot for liquid transfer. Specifically, tagmentation was done in 1.6ul (1.2ul Tagment enzyme mix, 0.4ul diluted cDNA) at 55°C, 10 min. Neutralization buffer was added 0.4ul per well and incubated at room temperature for 5 min to stop the reaction. Then 0.8ul Illumina 384 Indexes (0.4ul each, 5uM) and 1.2ul PCR master mix were added to amplify whole transcriptomes using the following program: (1) 72°C 3 min; (2) 95°C 30 sec; (3) 10 cycles of 95°C 10 sec, 55°C 30 sec, 72°C 1 min; (4) 72°C 5 min. Libraries from a single 384 plate were pooled together in an Eppendorf tube and purified twice with PCRClean DX beads. The quality and concentrations of the final mixed libraries were measured with Bioanalyzer and Qubit, respectively, before Illumina Nextseq sequencing.

For bulk RNA-seq, TMEM119^+^ cells from P60 brain regions were sorted (SONY SH800) into a 1.5ml Eppendorf tube containing 350ul RLT buffer (3000 cells per sample). RNA was extracted using RNeasy Micro Kit (Qiagen 74004) and eluted with 14ul nuclease-free water. To make cDNA, Smart-Seq v4 ultra low input RNA kit (Clontech 634890) was used following manufacture’s instruction (9.5ul RNA input). cDNA was amplified for 9 cycles, and cDNA concentrations were measured with Qubit. Libraries were prepared using Nextera XT DNA Sample Prep Kit (380pg cDNA per sample) following the standard protocol. Nextera 24 Indexes (Illumina 15055293) were used during whole transcriptomes amplification (13 cycles). Library quality was assessed with Bioanalyzer and Qubit. Nextseq sequencing was performed to a depth of (1.6±0.33) X10^7^raw reads per sample and (1.07±0.24) X10^7^ mapped reads per sample. This led to the detection of 11727±563 genes, comparable to 11407±2177 genes detected in P60 microglia from our previously published dataset (Bennett et al., 2016).

### Processing of scRNA-seq raw data and generation of gene counts

Prinseq was first used to filter sequencing reads shorter than 30 bp (-min_len 30), trim the first 10 bp at the 5’-end (-trim_left 10) of the reads, trim reads with low quality from the 3’-end (-trim_qual_right 25) and remove low complexity reads (-lc_method entropy, - lc_threshold 65). Then, Trim Galore was applied to trimmed the Nextera adapters (-- stringency 1). The remaining reads were aligned to the mm10 genome by calling STAR with the following options: --outFilterType BySJout, --outFilterMultimapNmax 20, -- alignSJoverhangMin 8, --alignSJDBoverhangMin 1, --outFilterMismatchNmax 999, - - outFilterMismatchNoverLmax 0.04, --alignIntronMin 20, --alignIntronMax 1000000, -- alignMatesGapMax 1000000, --outSAMstrandField intronMotif. Picard was then used to remove the duplicate reads (VALIDATION_STRINGENCY=LENIENT, REMOVE_DUPLICATES=true). Finally, the aligned reads were converted to counts for each gene by using HTSeq (-m intersection-nonempty, -s no).

### Quality control for scRNA-seq data

To filter out cells with low sequencing quality, three criteria, namely the number of total reads, the number of total detected genes and correlation coefficient between ERCC spike-ins input and corresponding read counts, were each evaluated based on the distributions of the data. The distribution of the total reads (in logarithmic scale) was fitted by a truncated Cauchy distribution, and data points in two tails of the estimated distribution were considered as outliers and eliminated. Fitting and elimination were then applied to the remaining data. This process was run iteratively until the estimated distribution became stable. The threshold was set to the value where the cumulative distribution function of the estimated distribution reaches 0.05. Similarly, cells with small numbers of detected genes, and poor correlation coefficients for ERCC (low sequencing accuracy) were dropped. After filtering, 1816 cells, out of 1922, were retained.

### Sensitivity and accuracy assessment of scRNA-seq data using ERCC spike-ins

We used a previously published algorithm to estimate sensitivity and accuracy of scRNA-seq data based on ERCC spike-ins with known concentrations (Svensson et al., 2017). To estimate sensitivity, for each cell, binomial logistic regression was used to fit the actual numbers of ERCC spike-in molecules and the responses (detected or not) and then the number of ERCC molecules (detection limit) corresponding to 0.5 detection probability was determined. Distribution of detection limit for all cells that passed quality control (QC) was plotted in Figure S1E, which indicates the overall sensitivity level of the assay. To estimate accuracy, Pearson correlation coefficient between the actual numbers of spike-in molecules used as input and the corresponding average reading counts across all cells (passed QC) was calculated and shown in Figure S1F.

### Clustering analysis of scRNA-seq data

The Seurat package was used to perform unsupervised clustering analysis on scRNA-seq data (Macosko et al., 2015). Briefly, gene counts for cells that passed QC were normalized to the total expression and log-transformed, and then highly variable genes were detected (y.cutoff=0.5). Depending on the analysis, cell cycle effect could be regressed out using the ScaleData function. Using highly variable genes as input, principal component analysis was performed on the scaled data in order to reduce dimensionality. Statistically significant principal components were determined by using the JackStrawPlot function. These principal components were used to compute the distance metric, which then generated cell clusters. Non-linear dimensional reduction (tSNE) was used to visualize clustering results. Differentially expressed genes were found using the FindAllMarkers (or FindMarkers) function that ran Wilcoxon rank sum tests.

### Differential gene expression analysis on region-specific microglia

To identify differentially expressed (DE) genes of adult microglia in each of the four regions (cerebellum, hippocampus, striatum, and cortex), we performed DE gene analysis (one versus rest) on both scRNA-seq and bulk RNA-seq data by using EdgeR. For scRNA-seq data, genes expressed (CPM>2) in at least 5% of total cells were tested by the likelihood ratio method for DE gene identification. For bulk RNA-seq data, genes that were detected (CPM>2) in at least 2 samples were used to perform quasi-likelihood F-test to identify DE genes across regions.

### Cell cycle regression on scRNA-seq data

We used Seurat package to regress out cell cycle effects in scRNA-seq data when analyzing P7 microglia heterogeneity. Briefly, each cell was assigned with a G2M score and an S score, based on its expression of G2/M and S phase markers. Then for each gene, the relationship between the gene expression and the G2M and S scores across all cells was modeled. A “corrected” expression matrix, determined by the scaled residuals of this model was obtained and used for downstream analysis.

### Pseudotime analysis of dividing microglia to generate phase-specific gene sets

This algorithm is based on a published method with modifications (Macosko et al., 2015). The same analysis was conducted on P7-C0 and P7-C1 microglia separately. Simply, 5 conserved gene sets from studies on HeLa cell lines that are periodically expressed in 5 cell cycle phases (G1/S, S, G2/M, M and M/G1) were curated (see below) and used as the initial seeds in order to discover microglia-specific cell cycle genes. Cells that had significantly different expression levels (ANOVA test) for the 5 curated gene sets were selected as “dividing cells” for pseudotime ordering, and the rest were temporarily labeled as “non-dividing cells”. Dividing cells were ordered based on expression of these phase-specific genes as previously described (Macosko et al., 2015). Then new genes that were expressed by cells non-randomly distributed along the order and correlated with average expression of gene seeds specific to any one or two adjacent phases, were assigned with a phase label and added to the corresponding seeds gene list (see below). After this, using updated seeds as input, the whole process was run iteratively, that is “cell selection”-“dividing cell ordering”-“seeds updating with new genes”, until the cell ordering was stable (the correlation between current ordering and previous ordering larger than 0.9) for more than ten times. During the iteration, the initial gene seeds were always kept in the updated list for the next round to avoid random drifting by noise. Once cell ordering became stable, genes that were frequently identified (larger than five times) in the last ten iterations were saved as final cell cycle phase-specific genes. The final gene lists were then used again to determine dividing cells versus G0 cells (cells expressing genes of all phases at high levels were manually excluded), and the final pseudotime order of dividing cells.

#### Initial gene seeds curation

Five sets of conserved genes, each specific to a single phase of the cell cycle (G1/S, S, G2/M, M and M/G1), were selected from a published dataset (Macosko et al., 2015). A gene with low correlation (R<0.3) between its expression level (log2(FPKM+1)) and the average expression of its own gene set was discarded from the list. To ensure the phase label for each remaining gene is applicable to microglia, correlations between its expression and the average expression of each gene set were calculated, and the gene was assigned to the gene set that had the maximal correlation with it. The correlation examination and re-assignment were run iteratively until no more genes were re-assigned. Such 5 gene sets were used as initial gene seeds.

#### New phase-specific gene discovery to update seeds

For each gene, a sliding window with a certain size (length of ordered cells/10) was used to find out the windows with maximal average expression and minimal average expression along the cell order. The difference between the maximal and minimal average expression was calculated as the observed difference. Then the order of the cells was randomly permutated, and the same sliding window operation was performed to obtain a permutation difference. Permutation was repeated for 1.0e+05 times to generate a distribution of permutation differences, with which the observed difference was compared to calculate a *p*-value. Genes with adjusted *p*-values less than 0.05 (non-randomly expressed along the cell order) were retained. To determine which phase(s) an identified gene belongs to, 10 scenarios of phase assignment were considered: five single phases and five dual phases (any two consecutive phases). Correlations between the gene expression and average expression of gene seeds corresponding to a single phase or two consecutive phases were calculated, and the phase(s) yielding the maximal correlation was assigned to the gene. Genes were ignored if they had poor correlations (R<0.3) with all of the ten scenarios mentioned above.

### Gene network construction for P7 microglia

A gene network was built to reveal the relationship between different genes in P7 cells. To this end, genes that contributed most to cellular heterogeneity were chosen and correlations between these genes were shown on a network graph with features explained below. Specifically, genes that were expressed in at least 3% of total cells and contributed to the first 15 principal components underlying sources of variances were used for analysis. After quantile normalization for each gene’s expression, graphical lasso was used to explore the relationship between these genes. Graphical lasso estimated a sparse precision matrix (i.e., inverse covariance matrix) of the variables (genes) by using a lasso (L1) penalty. Such precision matrix revealed the conditional relationships between genes, upon which a gene network was built. In the network, each vertex was a gene, and the weight of each edge was the absolute value of the corresponding entry in the precision matrix. If two genes were nonadjacent (value of the corresponding entry in the precision matrix was 0), they were conditionally independent, given all the other genes. For better visualization, the width of each edge in the graph was proportional to its weight. Edges with small weights were not shown to simply the illustration. If the value of an entry in the estimated covariance matrix was positive, the corresponding edge was a solid line, indicating a positive correlation between the two genes. A negative value for an entry was translated to a dashed line, indicating a negative correlation. The area of each vertex (gene) was proportional to the total weights of all edges connecting to it. To illustrate how genes in the network contributed to cell clustering, genes were painted with different colors based on whether they were differentially up-regulated in any P7 cluster and, if so, in which cluster(s). Genes shown in the network that were significantly up-regulated in a single P7 cluster were colored according to a predefined colormap (P7-C0=red; P7-C1=blue; P7-C2=green). Genes significantly up-regulated in two clusters were mapped to a new colormap, which included mixed colors of the two corresponding clusters (e.g. P7-C1&P7-C2= turquoise). The shades and tints of colors were determined by the logarithmic transform of adjusted *p*-values. Genes not differentially expressed were in grey.

### Developmental pseudotime analysis for microglia

Sequencing reads for 648 P7 microglia (excluding P7-C2), 218 P60 homeostatic microglia (excluding 3 cells with detectable *Clec7a* expression) and 133 P7-GPNMB^+^CLEC7A^+^ microglia were analyzed together to build developmental pseudotime using the Monocle package (Trapnell et al., 2014). First, clustering analysis was performed with Seurat after cell cycle regression as described above. Then the Seurat object was imported into Monocle, which used statistical models to find out differentially expressed (DE) genes based on the clustering result. Top 350 DE genes (*q*<0.01) were selected for cell ordering. Monocle then used reversed graph embedding (DDRTree method) to do dimensionality reduction. Finally, Monocle performed manifold learning in order to generate a tree-like structure reflecting the developmental trajectories from one cell state to others.

### GO term analysis

PANTHER Classification System was used to perform Gene Ontology term analysis (Mi et al., 2010). The input included 782 genes up-regulated in P7-GPNMB^+^CLEC7A^+^ microglia (compared with P7-C0 immature microglia), and statistical overrepresentation test was run using default settings. Fold enrichment values and false discovery rate (- log10 FDR) for the enriched GO-Slim Biological Process (BP) terms and GO-Slim Cellular Component terms were plotted. FDR was calculated by Fisher’s Exact with FDR multiple test correction.

### RNA *in situ* (RNAscope)

Mouse brains were rapidly dissected out and immediately embedded in OCT (optimal cutting temperature) compound (Tissue-Tek). The fresh frozen sections (sagittal) were prepared with cryostat at 12um thickness and RNA *in situ* was performed using RNAscope technology (Advanced Cell Diagnostics) following the manufacturer’s protocol (RNAscope Multiplex Fluorescent Assay). Briefly, after fixation and dehydration, the slides were treated with protease IV for 20 min at room temperature. RNAscope probes were hybridized for 2 hours at 40 °C and then fluorescence staining was done through a signal amplification system. Pre-designed Probes (ACD) against the following mRNA were used: *Cx3cr1 (314221), Gpnmb (489511), Spp1 (435191), Mbp (451491), Igf1 (443901), H2-Eb1 (509081), Lilra5 (514461), Itgax (311501), Tmem119 (472901), Fos (316921), Egr1 (423371)*.

### Immunohistochemistry

Mice were transcardially perfused with PBS followed with 4% para-formaldehyde (PFA). Dissected brains were fixed with PFA overnight at 4°C and then cryopreserved with 30% sucrose for 24 hours at 4°C. Tissues were embedded in the optimal cutting temperature (OCT) compound (Tissue-Tek). Free floating sagittal sections (50um) were stored in 6- well plates with PBS and then blocked and permeabilized with 10% serum/0.2% Triton-X in PBS for 15 min at room temperature. Sections were incubated with primary antibodies at 4°C overnight followed by secondary antibody staining for 2 hours at room temperature. After washes, the sections were transferred onto glass slides and mounted in Vectashield with DAPI (Vector Laboratories H-1200). Images were acquired using Zeiss LSM 880 inverted confocal and Zeiss Axio Imager fluorescence microscopy, and imported into Image J for quantification or Imaris for 3D reconstruction.

The following primary antibodies were used: Goat polyclonal anti IBA1 (Abcam ab5076, 1:500); Rabbit monoclonal anti TMEM119 (clone 28-3) (Abcam ab209064, 1:100); Sheep polyclonal anti c-FOS (Abcam ab6167, 1:200); Rabbit monoclonal anti EGR1 (clone 15F7) (Cell Signaling Technology 4153, 1:50); Rabbit polyclonal anti cleaved CASPASE-3 (Asp175) (Cell Signaling Technology 9661, 1:200); Rat neutralizing monoclonal anti-mDECTIN-1-IgG (clone R1-4E4) (Invivogen mabg-mdect, 1:30); Rat monoclonal anti Myelin Basic Protein (clone 12) (Abcam ab7349, 1:100); Rat monoclonal anti-DECTIN 2 (clone D2.11E4) (Abcam ab34724, 1:100); Rabbit monoclonal anti PDGF Receptor α (D1E1E) (Cell Signaling Technology 3174S, 1:500); Goat polyclonal anti Osteopontin/OPN (R&D Systems AF808, 1:13). The following secondary antibodies were used: Donkey anti-goat IgG (H+L) cross-adsorbed, Alexa Fluor 488 (Thermo Fisher Scientific 11055, 1:1000); Donkey anti-rabbit IgG (H+L), Alexa Fluor 488 (Thermo Fisher Scientific 21206, 1:1000); Donkey anti-sheep DyLight 594 (Jackson Immuno Research Laboratories 713-586-147, 1:200); Donkey anti-rabbit IgG (H+L) Highly cross adsorbed, Alexa Fluor 594 (Thermo Fisher Scientific 21207, 1:200); Donkey anti-rat IgG (H+L) highly cross-adsorbed, Alexa Fluor 594 (Thermo Fisher Scientific 21209, 1:200); Donkey anti-rabbit IgG (H+L) highly cross-adsorbed, Alexa Fluor 647 (Thermo Fisher Scientific 31573, 1:200); Donkey anti-rat IgG (H+L) AffiniPure, Alexa Fluor 647 (Jackson Immuno Research Laboratories 712-605-153, 1:200).

### Phagocytosis assay by slice culturing system

P7 mice were decapitated and dissected brains were immediately put in pre-cooled culture medium with serum (65% MEM (Sigma #M2279); 10% horse serum; 25% HBSS; 6.5 mg/ml glucose; 2mM Glutamine; 1% Pen/Strep), or serum-free microglial culture medium (DMEM/F12, 1% Pen/Strep; 2 mM glutamine; 5 ug/mL N-acetyl cysteine; 5 ug/mL insulin; 100 ug/mL apo-transferrin; 100 ng/mL sodium selenite; 2 ng/mL human TGF-2; 100 ng/mL murine IL-34; 1.5 ug/mL ovine wool cholesterol; 10 ug/mL heparan sulfate; 0.1 ug/ml oleic acid; 0.001 ug/ml gondoic acid) (Bohlen et al., 2017). The entire procedure was done on ice with pre-cooled solutions until culturing. Sagittal sections were prepared with vibratome (Leica VT1000S) at 250um thickness and then transferred to insert wells (Millicell Cell Culture Insert, 30 mm, Millipore #PICM03050) on a 6-well plate with medium. The plate with the sections was incubated for 1 hour in the incubator (37°C, 5% CO2) before pHrodo™ Red Zymosan Bioparticles (Thermo Fisher Scientific P35364) were added at 0.5mg/ml to cover the whole sections (about 150ul each). Then the plate was incubated for 4 hours for phagocytosis (37°C, 5% CO2). After washes with PBS, sections were fixed with 4% PFA for 30 min at room temperature and then blocked with 10% serum/0.2% Triton-X in PBS for 15 min at room temperature. Immunohistochemistry was performed as described above.

### 3D reconstruction of confocal images

Confocal z-stacks were imported into Imaris software (Bitplane AG, Zurich, Switzerland). Regions of interest were isolated to make IsoSurfaces for each channel. To quantify interactions between postnatal WAM and other neural cell types, whole sections were 3D reconstructed, and overlapping volumes between postnatal WAM and corresponding cell types were obtained with the function of distance transformation. The percentage of the overlapping volume over the volume of a given cell (MPB^+^, Aldh1l1-GFP^+^ or PDGFRA^+^) was calculated to represent the level of interaction between this cell and the postnatal WAM. Substantial interaction was defined as 30% overlapping volume in the cell. Similar results were observed with higher percentages.

